# Wild animal oral microbiomes reflect the history of human antibiotics use

**DOI:** 10.1101/2020.12.22.423960

**Authors:** Jaelle C. Brealey, Henrique G. Leitão, Thijs Hofstede, Daniela C. Kalthoff, Katerina Guschanski

**Author notes:** Department of Natural History, NTNU University Museum, Trondheim, Norway.

## Abstract

Following the advent of industrial-scale antibiotics production in the 1940s, antimicrobial resistance (AMR) has been on the rise and now poses a major global health threat. Because AMR can be exchanged between humans, livestock and wildlife, evaluating the potential of wild animals to act as AMR reservoirs is essential. We used shotgun metagenomics sequencing of dental calculus, the calcified form of the oral microbial biofilm, to determine the abundance and repertoire of AMR genes in the oral microbiome of Swedish brown bears from museum specimens collected over the last 200 years. Our temporal metagenomics approach allowed us to establish a baseline of natural AMR in the pre-antibiotics era and to quantify a significant increase in total AMR load and diversity of AMR genes that is correlated with human antibiotics use. We also demonstrated that Swedish public health policies were effective in limiting AMR spillover into wildlife.

## Introduction

Since the discovery and use of penicillin to treat bacterial infections in the 1930s and 1940s, the hopes and promises of new antibiotics have been curbed by the rapid rise of resistance to each new class of drugs (Landecker 2016). This trend has been exacerbated by the misuse and overuse of antibiotics in both the human healthcare and agriculture sectors (Levy and Marshall 2004; Landers et al. 2012). Antimicrobial resistance (AMR) now poses a serious global public health threat in terms of mortality, morbidity and economic burden (Naylor et al. 2018; Thorpe et al. 2018). In Europe alone, resistant bacterial infections account for approximately 33,000 deaths and 875,000 disability-adjusted life-years per year (Cassini et al. 2019). Globally, 10 millions of death per year are predicted from resistant infections by 2050 (O’Neill (chair) 2014).

The challenges of combating AMR are partly due to the wide diversity of antibiotic resistance genes (ARGs) that are present in (putatively pathogenic) microorganisms. Some ARGs confer resistance to a broad range of antibiotics, such as the various families of multidrug efflux pumps. Others target a specific class of antibiotics, such as the beta-lactamases, which inactivate beta-lactam antibiotics like penicillin (Alekshun and Levy 2007). In addition, ARGs are readily shared between different bacterial taxa within complex microbial communities via horizontal gene transfer primarily of mobile elements, such as plasmids (Thomas and Nielsen 2005). Thus, microbial communities, including those residing within human or animal hosts (host microbiomes), can harbour a diverse repertoire of ARGs that can be transmitted between commensals and putative pathogens (Salyers et al. 2004; Arnold et al. 2016). Exposure to antibiotics subjects the host microbiome to strong selective pressure that favours resistant microorganisms and reduces or eliminates susceptible strains, consequently leading to the perturbation of the entire microbial community (Francino 2016).

It is becoming increasingly clear that wild and domestic animals are reservoirs for potential human pathogens, including resistant bacteria (Jones et al. 2008). The One Health approach recognises the co-dependence of human health, animal health and the health of ecosystems and the environment as a whole (Gibbs 2014). In the context of AMR, the spillover of antibiotics and resistant bacteria from human use into the environment can re-enter human populations from environmental and wildlife reservoirs. Antibiotics and resistant bacteria from human production and use in both clinical and agricultural settings can leak into the environment through waste production (Miller et al. 2009; Pruden et al. 2012; He et al. 2016; Zhu et al. 2017). Wildlife can then come into contact with both antibiotics and resistant bacteria by means of contaminated soil, water and food sources (Mariano et al. 2009). Resistant bacteria can also be transmitted between wild animals and humans directly (e.g. during hunting, trapping, treatment at wildlife rehabilitation centres) (Jijón et al. 2007; Zottola et al. 2013; Mo et al. 2018) and between wild and domesticated animals through direct contact and exposure to waste products (e.g. rodents and flies residing on livestock farms) (Leatherbarrow et al. 2007; Kozak et al. 2009; Literak et al. 2009; Luque et al. 2009; Guenther et al. 2010; Dolejska et al. 2011; Navarro-Gonzalez et al. 2012; Carlson et al. 2015; Nhung et al. 2015). While several studies have provided evidence for the exchange of ARGs between humans, livestock and wildlife (Leatherbarrow et al. 2007; Literak et al. 2009; Dolejska et al. 2011; Subbiah et al. 2020), the impact of human usage of antibiotics on the environment in general and wildlife populations in particular remains little explored (Goulas et al. 2020).

One of the difficulties in quantifying the impact of human antibiotic usage on the environment is the fact that AMR is ubiquitous in nature. Some bacteria and fungi naturally produce antibiotics (Martin and Liras 1989) and AMR has thus evolved as protection against both self-generated antibiotics (Tahlan et al. 2007) and those produced by competing species (Hibbing et al. 2010). Some genes that convey AMR have been co-opted for their general detoxifying function. For instance, the multidrug efflux pumps eject both antibiotics and other toxic compounds from the bacterial cell, such as heavy metals, detergents and host-derived molecules like bile salts (Poole 2005). ARGs have been found across the globe, including in the most pristine environments free from human activities (Miller et al. 2009; Bhullar et al. 2012; Nesme et al. 2014; Goethem et al. 2018). It is therefore difficult in many cases to establish a natural, human-unaffected AMR baseline and to distinguish between human-associated and natural sources of AMR (Allen et al. 2010; Vittecoq et al. 2016).

Time can serve as a substitute for space in the study of AMR. Collections of bacteria isolates from the ‘pre-antibiotic era’ in the early 20th century have shown resistance to modern antibiotics (Smith 1967; Hughes and Datta 1983; Fusté et al. 2012). Recent advances in the field of ancient DNA have also allowed for the characterisation of ARGs from time periods that pre-date human antibiotics production, which reflect the natural AMR potential of a given environment. Diverse ARGs have been reported from culture-based and metagenomic studies of permafrost cores ranging from thousands to tens of thousands of years in age (Dcosta et al. 2011; Perron et al. 2015; Filippova et al. 2019; Okubo et al. 2019). ARGs have also been detected in host-associated microbiomes from specimens collected from before the industrial-scale production of antibiotics, in particular from dental calculus samples. Dental calculus is the calcified form of the dental plaque microbial biofilm that forms on mammalian teeth (Jin and Yip 2002). It is built up periodically throughout an individual’s life and preserves DNA and proteins from the oral microbiome within a calcified matrix, protected from invasion from external microorganisms following the host’s death (Adler et al. 2013; Warinner et al. 2014). Metagenomic sequencing of dental calculus samples has detected ARGs in medieval humans (Warinner et al. 2014) and 19-20th century wild animals (Brealey et al. 2020), as well as 15th century human paleo-faeces (Rifkin et al. 2020). With appropriate sampling, it is therefore possible to use dental calculus as a source of host-associated microbiomes that could reflect the history of AMR through time.

Here, we studied the progression of AMR through time in host-associated microbiomes of wild brown bears (*Ursus arctos*) in Sweden by characterising the abundance and repertoire of ARGs from bear dental calculus. We have previously shown that metagenomic sequences from dental calculus are a rich source of information on the oral microbial community of brown bears, including potential pathogens and ARGs (Brealey et al. 2020). As wide-ranging omnivores and scavengers, brown bears have a diverse diet and are exposed to a variety of potential sources of AMR from both prey species and the environment (Dahle et al. 1998; Elfström et al. 2014; Vittecoq et al. 2016). While they are generally solitary and prefer remote areas, Swedish brown bears do approach human settlements, predate on livestock and occasionally consume crops (Dahle et al. 1998; Dahle and Swenson 2003; Elfström et al. 2014). Direct human contact can also occur, primarily with hunters and their hunting dogs (hunting either bears or their prey species, e.g. moose), as well as less frequent or indirect contact with humans and pets through recreational hiking, berry-picking and forestry work (Støen et al. 2018). We used dental calculus from museum-preserved Swedish brown bear specimens that span the last 200 years and thus partly predate the industrial-scale production of antibiotics that started in Sweden in the 1950s (Wickman 1969). The temporal sampling allowed us to determine the baseline ARGs of the naïve brown bear oral microbiome from before the 1950s and to quantify changes in prevalence and diversity of ARGs in the following decades.

Sweden has a well-documented history of antibiotic use and control in both humans and domestic animals. While industrial-scale production of antibiotics started in the mid-1940s in the US (Landecker 2016), antibiotics were not commercially produced in Sweden until after 1947 (Wickman 1969). Antibiotic production and usage increased in Sweden in the 1950s and 1960s, including the use of antibiotics at lose doses in livestock to improve their growth and feed efficiency (antibiotic growth promoters) (Wickman 1969; Begemann et al. 2018). Antibiotic usage reached its peak in the 1970s and 1980s, when increased concerns about mounting antibiotic resistance resulted in voluntary decreases in agricultural use (Edqvist and Pedersen 2001). In 1986, Sweden banned the use of antibiotic growth promoters in agriculture (Wierup 2001). In 1995, the Swedish strategic program against antibiotic resistance (Strama) was founded in response to a rapid increase in penicillin resistance, particularly among young children (Ekdahl et al. 1998; Folkhälsomyndigheten 2014). Since then, Sweden has implemented a number of long-term measures to promote the rational use of antibiotics, regulate the sales of antibiotics for humans and animals and continuously monitor AMR (Folkhälsomyndigheten 2014). Consequently, sales of antibiotics for both outpatient care and veterinary use have generally decreased (Folkhälsomyndigheten and SVA 2019). As of 2018, overall AMR in humans and animals in Sweden is low from an international perspective (Folkhälsomyndigheten and SVA 2019). We therefore used our dental calculus samples to investigate the effect of both antibiotic overuse in the 1970s and 1980s and subsequent antibiotic control strategies implemented by Swedish public health authorities in the 1990s on ARG abundance and diversity in Swedish brown bears. We hypothesised that increased human use of antibiotics led to increased prevalence and diversity of ARGs in wildlife microbiomes. We further predicted that proximity to human habitation increased exposure to human waste products and thus resulted in higher AMR load in individuals with home ranges in more densily populated areas.

## Results

### Sample processing and authentication

We sequenced 82 dental calculus samples collected from healthy teeth of Swedish brown bear specimens from across central and northern Sweden dating between 1842 and 2016 (Figure 1). After quality control and read processing (including filtering of host-derived reads, see Methods), each calculus sample contained on average 16 million reads (range 34 reads to 186 million reads). Following microbial taxonomic assignment with Kraken2 (Wood and Salzberg 2014) and species-level abundance estimation with Bracken (Lu et al. 2017), three calculus samples were excluded due to low microbial read content (see Methods).

**Figure 1.**
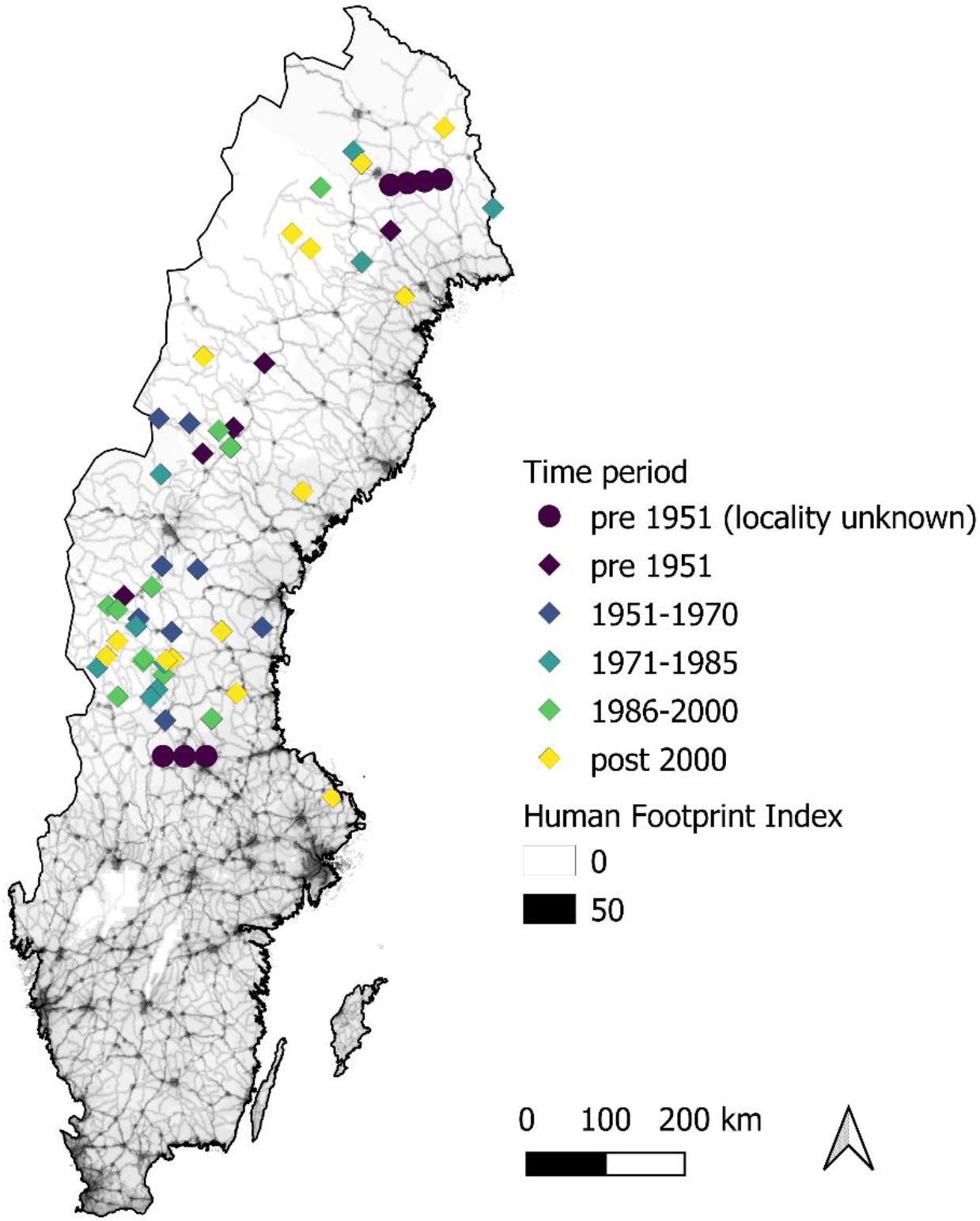
Sampling locations of Swedish brown bear museum specimens included in this study (n=57 following filtering and processing steps, see Methods). Specimens are coloured by the time period in which they were collected. Samples are plotted according to their GPS coordinates, locality location information as obtained from museum records (most specimens collected before 1995) or arbitrary coordinates within their counties, when only this level of geographic information was available (seven samples from pre-1951, shown by circles). The underlying terrain of Sweden is shaded by the 2009 Human Footprint Index (increased human activities/populations corresponds to darker shaded areas). The map of Sweden was obtained from Lantmäteriet (https://www.lantmateriet.se/sv/Kartor-och-geografisk-information/geodataprodukter/produktlista/oversiktskartan) and the Human Footprint Index data from NASA Socioeconomic Data and Applications Center (https://doi.org/10.7927/H46T0JQ4).

While endogenous microbial DNA is generally well-preserved in mammalian dental calculus, samples can be contaminated by environmental DNA from museum storage and laboratory processing (Key et al. 2017; Mann et al. 2018). Thus, to control for background museum and laboratory contamination, we sequenced two swabs taken from the museum storage facility (see Methods), 29 extraction blank samples and 8 library preparation blank samples. Microbial source analysis with the Bayesian classification tool SourceTracker (Knights et al. 2011) demonstrated that the majority of the calculus samples included microbial taxa also found in human dental plaque and calculus microbial communities, whereas blanks and swabs were generally more similar to human skin and soil environments (Supplementary Figure S1). In an ordination of the microbial taxa abundances, extraction and library preparation blanks formed a tight cluster, whereas the more variable bear calculus samples tended to be separated by the proportion of oral microbiome taxa (human plaque + calculus SourceTracker estimates; Supplementary Figure S2 and Supplementary Table S1). We therefore excluded 22 bear calculus samples with < 5% of the microbiome community similar to the human oral microbiome, retaining 57 samples for further analyses.

To minimise the possibility that modern environmental resistant bacteria may confound our estimates of AMR in the endogenous bear oral microbiome, we restricted our analysis to the subset of bacteria that are known to colonise human and pet oral cavities and are thus unlikely to result from environmental contamination (Supplementary Table S2). All blank samples had less than 250 reads assigned to oral bacteria, whereas the two swabs had read counts that were generally an order of magnitude lower than the 57 retained bear dental calculus samples (mean swabs = 44,162, mean samples = 1,135,500, Supplementary Figure S3). We then blasted the oral bacteria reads against the Comprehensive Antibiotic Resistance Database (CARD) (Jia et al. 2017). The top match for each read was assigned to its respective gene family under the Antibiotic Resistance Ontology (ARO). On average, 96 reads per bear sample were assigned a match in CARD (median: 48 reads, range: 0 – 497), whereas no oral bacteria reads from the blanks or swabs had a match in CARD.

### Total AMR load reflects antibiotics usage in Sweden

To determine how AMR prevalence has changed over time in brown bear oral microbial communities, we binned the bear samples into 5 time periods based on historical antibiotic usage in Sweden: those collected before 1951 (the pre-antibiotic era), those collected between 1951-1970 and 1971-1985 (reflecting increasing usage), those collected between 1985-2000 (when control measures were first implemented) and those collected after 2000 (post control measures).

We detected ARGs in samples collected in the pre-antibiotic era (Figure 2), in line with the expectation that AMR is a natural function of microbial communities independent of contribution from human use. Total AMR load (total number of reads with a match in CARD divided by the total number of oral bacteria reads in a sample) increased from the 1950s through 1990s with increasing use of antibiotics, before decreasing after the implementation of control measures in the 2000s (Figure 2). Both the increase in the decades before 2000 and the decrease post 2000 were statistically significant (generalised linear model with a quasibinomial distribution, Table 1). Among possible confounding variables, only median length of the oral bacteria DNA fragments was significantly correlated with total AMR load (Spearman correlation rho = 0.457, *p* = 0.00035; Supplementary Figure S4, Supplementary Table S3), suggesting that ARGs are more likely to be detected in samples with longer DNA fragments. However, the significant relationship between time period and total AMR load remained after controlling for median DNA fragment length (Table 1). Thus, we detected a strong temporal correspondence between total AMR load in bear dental calculus and human use of antibiotics.

**Table 1.**
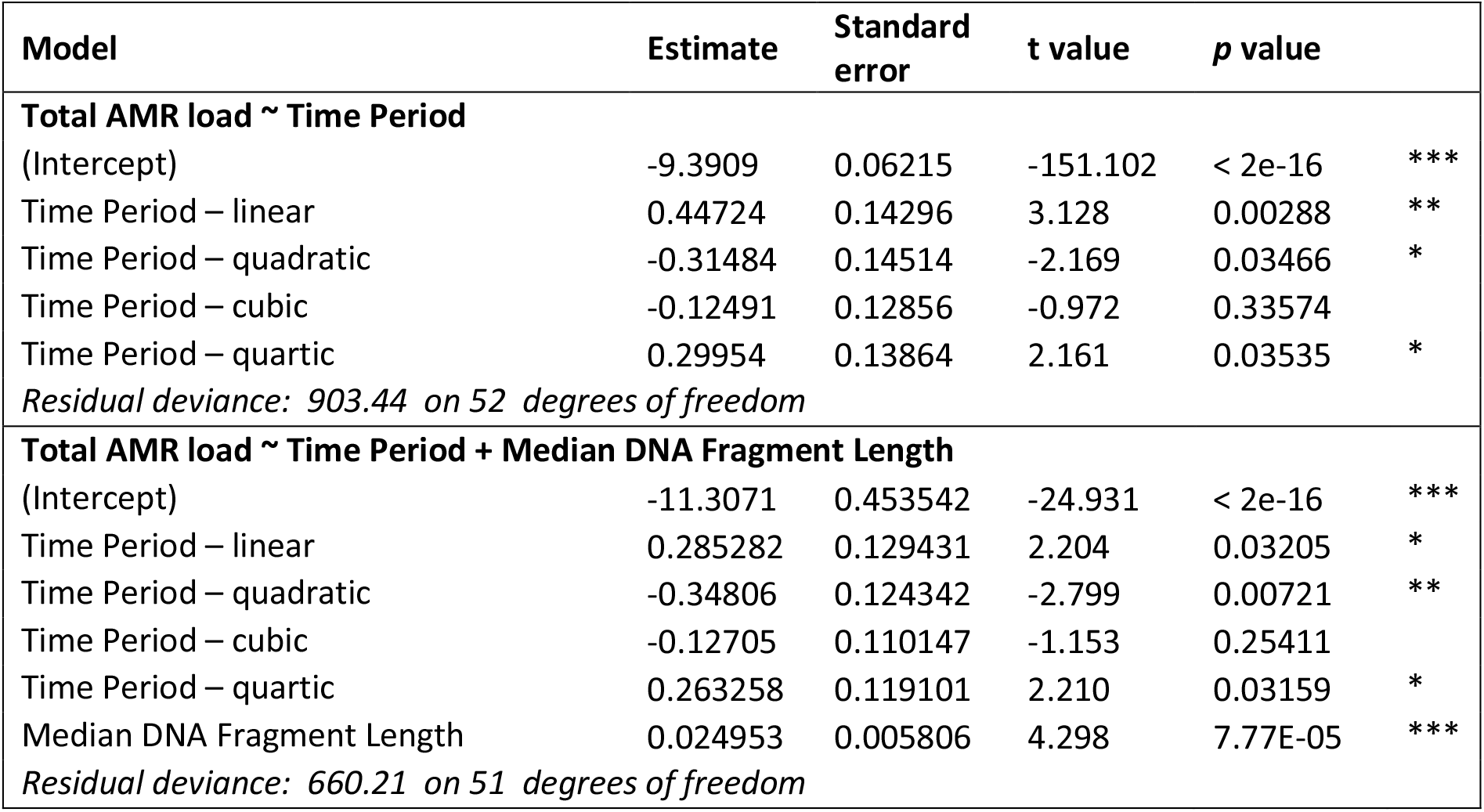
Results from generalised linear models of changes in total AMR load across time period, with and without controlling for median DNA fragment length. Time period was coded as an ordered factor with five levels, thus results are provided for each polynomial (linear through to quartic).

| Model | Estimate | Standard error | t value | p value |  |
| --- | --- | --- | --- | --- | --- |
| **Total AMR load ~ Time Period** |  |  |  |  |  |
| (Intercept) | -9.3909 | 0.06215 | -151.102 | < 2e-16 | *** |
| Time Period – linear | 0.44724 | 0.14296 | 3.128 | 0.00288 | ** |
| Time Period – quadratic | -0.31484 | 0.14514 | -2.169 | 0.03466 | * |
| Time Period – cubic | -0.12491 | 0.12856 | -0.972 | 0.33574 |  |
| Time Period – quartic | 0.29954 | 0.13864 | 2.161 | 0.03535 | * |
| *Residual deviance: 903.44 on 52 degrees of freedom* |  |  |  |  |  |
| **Total AMR load ~ Time Period + Median DNA Fragment Length** |  |  |  |  |  |
| (Intercept) | -11.3071 | 0.453542 | -24.931 | < 2e-16 | *** |
| Time Period – linear | 0.285282 | 0.129431 | 2.204 | 0.03205 | * |
| Time Period – quadratic | -0.34806 | 0.124342 | -2.799 | 0.00721 | ** |
| Time Period – cubic | -0.12705 | 0.110147 | -1.153 | 0.25411 |  |
| Time Period – quartic | 0.263258 | 0.119101 | 2.210 | 0.03159 | * |
| Median DNA Fragment Length | 0.024953 | 0.005806 | 4.298 | 7.77E-05 | *** |
| *Residual deviance: 660.21 on 51 degrees of freedom* |  |  |  |  |  |

**Figure 2.**
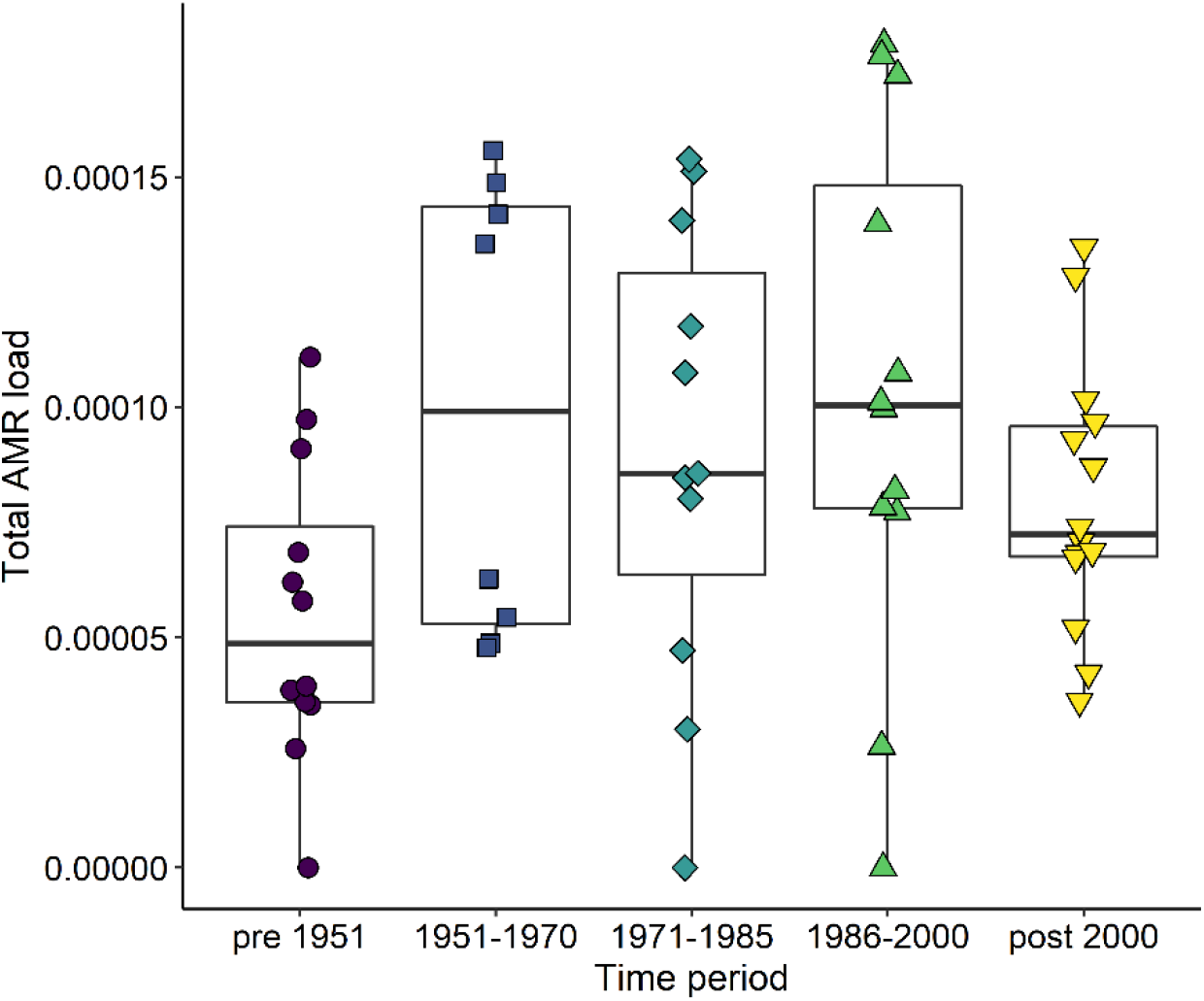
Total AMR load in bear calculus samples changes through time. Total AMR load was calculated as the number of reads mapping to CARD divided by the total number of oral bacterial reads in a sample. Each symbol within a given time period represents a unique brown bear dental calculus sample.

### ARG diversity increases over time

Next, we characterised and quantified ARG families in bear samples at different time periods to test if exposure to an increasing variety of antibiotics has had an impact on the diversity of ARGs present in wildlife host-associated microbiomes. The overall diversity of ARG families increased from 1986 onwards (Figure 3a), as reflected by the detection of novel rare ARGs in the latest two time periods (Figure 3b). This observation was not explained by differences in sample sizes (Figure 3a) or amount of data available (number of sequencing reads mapping to ARG families, Supplementary Figure S5) across time periods. Among these rare ARG families detected after 1985, we found multiple enzymes that are commonly encoded on plasmids and other mobile genetic elements. These included aminoglycoside modifying enzymes (ANT, APH, AAC; see Supplementary Table S4 for details) that target aminoglycosides like streptomycin, and beta-lactamases (OXA) that target beta-lactams like penicillin (Alekshun and Levy 2007). It is noteworthy that the detection of beta-lactamases temporally coinsides with the increase in use of narrow spectrum penicillins, which doubled in Sweden in the early 2000s (The Center for Disease Dynamics Economics & Policy 2018).

**Figure 3.**
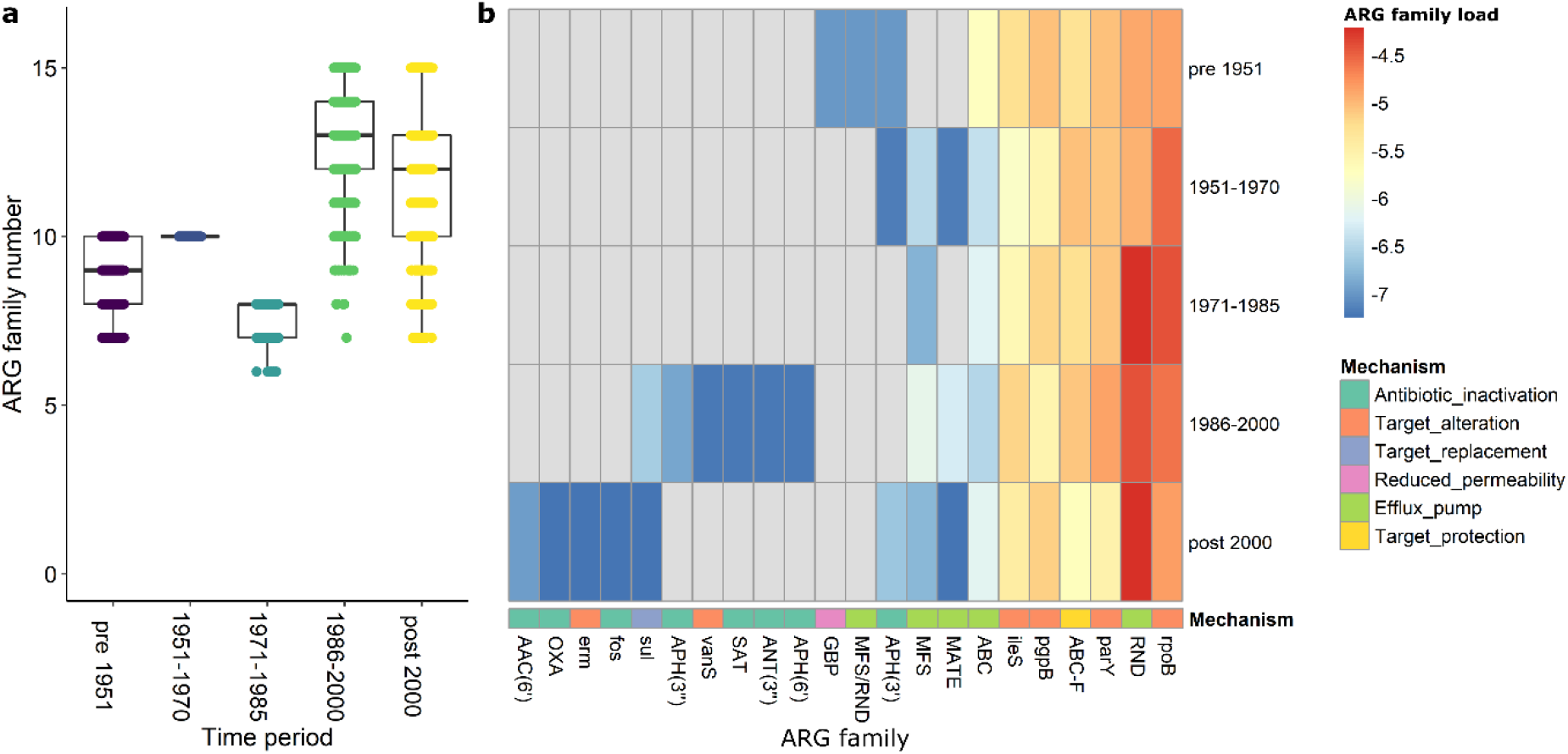
Diversity of ARG families changes over time. (a) Boxplots of the number of unique ARG families detected in each time period after subsampling to eight samples per time period (the lowest sample size available for time period 1951-1970) with 1000 independent repeats to control for differences in sample sizes between time periods. (b) Heatmap of log-transformed ARG family loads (proportion of oral bacteria reads mapping to each ARG family) in samples pooled by time period. ARG families that were not detected in a given time period are coloured grey. ARG families are annotated by their main mechanism of resistance recorded in CARD. ARG family abbreviations are described in Supplementary Table S4.

ARG families that were consistently detected in all time periods, including in the pre-antibiotic era, included multidrug efflux pumps, the ribosome-binding ABC-F family and ARGs conveying AMR through alterations in antibiotic targets, such as mutations in antibiotic binding sites or modification of cell wall components (Figure 3b). Multidrug efflux pumps, like the detected resistance-nodulation-cell division (RND) and ATP-binding cassette (ABC) efflux pumps, have a broad range of substrates, including multiple classes of antibiotics, solvents and toxic compounds (Poole 2005) and have many functions involved in bacterial colonisation and survival in the host environment (Jerse et al. 2003; Warner et al. 2008). ABC-F ribosomal protection proteins are closely related to ABC efflux pumps but show no efflux function. ABC-F proteins are common to all domains of life and bind to the ribosome, triggering the release of a wide range of antibiotics targeting the ribosome, thus rescuing inhibition of translation (Sharkey et al. 2016). The remaining commonly detected ARG families (rpoB, parY, ileS and pgpB) are usually encoded on the chromosome and confer resistance against specific antibiotic classes, such as rifamycin-resistant beta-unit of RNA polymerase (rpoB), aminocoumarin-resistant topoisomerase IV (parY) and mupirocin-resistant isoleucyl-tRNA synthetase (ileS). Lipid A phosphatase (pgpB) modifies an important component of the Gram-negative bacterial cell wall (lipid A in lipopolysaccharides), thus reducing susceptibility to peptide antibiotics like polymyxin B (Coats et al. 2009). Previous metagenomic studies of host-associated samples from the pre-antibiotics era have also detected multidrug efflux pumps, resistant rpoB and resistant DNA topoisomerase (Warinner et al. 2014; Rifkin et al. 2020). It is therefore unsurprising that these ARG families were detected in our pre-antibiotic era brown bear samples and adds further support to other studies finding that some classes of ARGs are ‘ancient’ (Dcosta et al. 2011).

### Changes in ARG abundances over time reflect changes in bacteria abundance

We observed changes through time in the abundance of the most frequent ARG families. Here, we focused on families that were detected throughout the study period. Total abundance of ABC-F proteins and resistant rpoB reflected the pattern of total AMR load, with an increase from 1951 to 2000 and a decrease post 2000 (Supplementary Table S5, Figure 4a-b). In contrast, RND efflux pump consistently increased over time to the present (Figure 4c). The other four most abundant ARG families (resistant ileS, ABC efflux pump, lipid A phosphatase and resistant parY) showed varied abundance trajectories over time. However, resistant ileS and lipid A phosphatase increased in load in the last two time periods, whereas parY showed higher abundance from 1950 to 2000, supporting the general pattern (Supplementary Table S5, Supplementary Figure S6).

**Figure 4.**
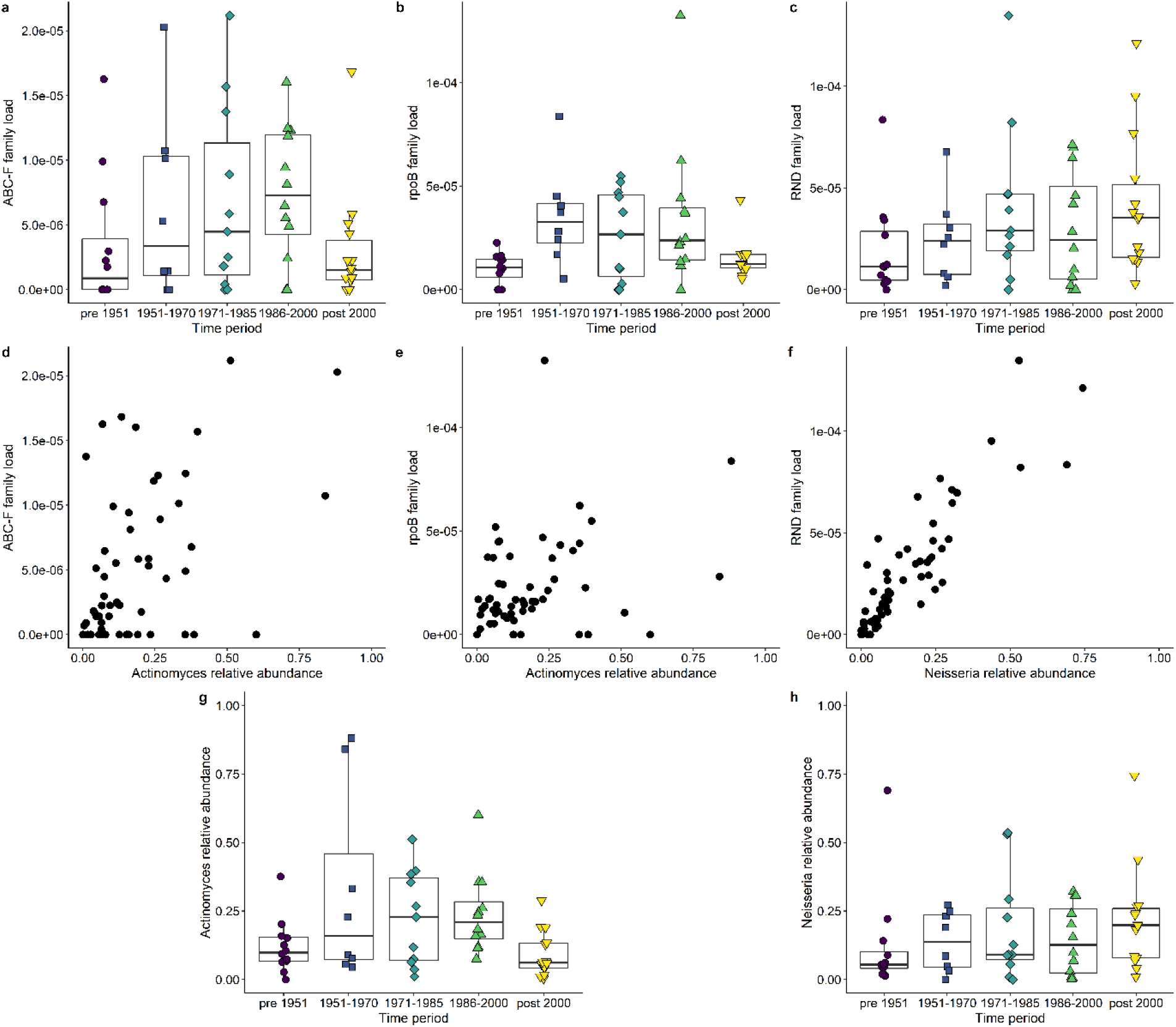
Changes in load of three of the most abundant ARG families in bear calculus are associated with changes in abundance of oral bacteria genera. (a-c) Proportion of oral bacteria reads mapping to (a) ABC-F (ATP-binding cassette) ribosomal protection protein, (b) antibiotic resistant rpoB (RNA polymerase beta subunit) and (c) RND (resistance-nodulation-cell division) antibiotic efflux pump across the study time periods. (d-f) Proportion of oral bacteria reads assigned by Bracken to *Actinomyces* is correlated with ABC-F (d) and antibiotic resistant rpoB (e) load, whereas proportion of reads assigned to *Neisseria* is correlated with RND efflux pump load (f). (g-h) Abundance of *Actinomyces* (g) and *Neisseria* (h) across the study time periods.

Changes in ABC-F and rpoB loads over time were correlated with changes in the abundance of the genus *Actinomyces* (Figure 4d-e,g) (ABC-F: similarity score = 0.443, *q* value = 0.0028; rpoB: similarity score = 0.401, *q* value = 0.0065). *Actinomyces* species are commonly found in the oral cavity, particularly as early colonisers of the tooth surface in the first steps of dental plaque formation, and thus some species are associated with dental caries infections in humans (Könönen and Wade 2015). However, not much is known about AMR potential of oral *Actinomyces* strains, since the genus rarely causes disease in humans and is usually susceptible to beta-lactams (Könönen and Wade 2015). RND abundance was strongly correlated with the abundance of the genus *Neisseria* (Figure 4f,h) (similarity score = 0.545, *q* value < 0.001). *Neisseria* is also known to colonise the oral cavity of mammals (Dent 1982; Dewhirst et al. 2012; Heydecke et al. 2013; Dewhirst et al. 2015) and has a well-characterised RND system (mtrCDE) which exports a wide variety of antibiotics, biocides and detergents (Chitsaz et al. 2019). Resistant ileS abundance was correlated with changes in the abundance of the genera *Parvimonas* and *Gemella* (*Parvimonas*: similarity score = 0.548, *q* value < 0.001; *Gemella*: similarity score = 0.363, *q* value = 0.0069), which both tended to show increased abundance in the last two time periods (Supplementary Figure S7).

Despite the detected changes in specific oral bacteria genera and the associated ARGs as described above, we did not observe changes through time in the overall oral bacteria community at the species-level (Supplementary Figure S8). Community composition was not associated with time period or total AMR load (Supplementary Table S1).

### Total AMR load is not associated with proximity to humans or geography

We hypothesised that bears collected from locations close to human habitations would have higher total AMR loads, as they might be more exposed to spillover of antibiotics or human-associated resistant bacteria. Fifty bear specimens had reliable location information (Figure 1), whereas the remaining seven had only the municipality or county recorded. We used historical records from Swedish parishes to estimate the human population density for the location of each brown bear sample (averaging across a county for the seven samples with imprecise locality information). Historical land use and human impact data were not available for the majority of sample localities. We instead used the 2009 Human Footprint Index (Figure 1), which combines data on human population density, land use, infrastructure and transport networks into a single measure, to estimate the magnitude of human impact in a 12.5 kilometre radius around the reported location of each of the 50 bears with known locations.

In contrast to our expectations, we found no association between total AMR load and historical human population density or modern Human Footprint Index (Supplementary Figure S9), nor with geographic regions, including municipality and county location (Supplementary Table S6). No association between geographic location and total AMR load was observed for any given time period (Supplementary Figure S10, Supplementary Table S6), suggesting a global change in total AMR through time and space in Sweden.

## Discussion

### AMR in wildlife host-associated microbiomes mimics human antibiotics use and production

Human production and use of antibiotics has been increasing since the 1940s (Landecker 2016). This trajectory has been followed by increases in AMR in human populations (Levy and Marshall 2004). It may therefore be expected that AMR has also increased in the environment and in domestic and wild animal populations as a result of spillover from human sources (Allen et al. 2010). Furthermore, development and mass use of new antibiotic classes may have diversified selective pressure on the naturally occurring AMR and led to a greater diversity of ARGs in more recent times. Here, we explicitly used temporal metagenomics of host-associated microbiomes to follow the progression of AMR in wild animals. This approach is best suited for studying changes in AMR and ARG diversity over historical time periods, as it minimises the difficulty in distinguishing between naïve and human-induced AMR. In accordance with expectations of spillover from human use, we found that the total AMR load and diversity of ARGs in the brown bear dental calculus microbiome closely follows the historically affirmed use of antibiotics in Sweden. More specifically, we detected increased total AMR load from the 1950s to 2000 and greater ARG diversity particularly since 1985, compared to the pre-industrial AMR background (before the 1950s).

Few other studies have investigated the temporal changes in AMR throughout the history of human antibiotics use. Increasing levels of AMR have been detected in archived soil samples from the 1940s to the mid-2000s (Knapp et al. 2010). Collections of human-associated bacteria isolated prior to 1955 have been found to have low levels of AMR (Smith 1967; Hughes and Datta 1983), although other studies utilising such collections have found similar AMR profiles in both pre-1950s and modern strains (Fusté et al. 2012). In wildlife, changes in AMR levels have only been investigated over the last 20 years, where AMR was found to increase from 2004 to 2010 in bacteria isolated from stranded marine mammals (Wallace et al. 2013). Our study thus represents the first systematic quantification of AMR in wild animals over the course of the last 200 years.

While the spillover of human-produced antibiotics and resistant bacteria into natural systems has been well documented, our results suggest that this process may be reversible. Sweden was one of the first countries to impose legislation to control the use of antibiotics in agriculture in 1985 and today has a strong antibiotic stewardship program with thorough monitoring of resistance in human and animal infections and screens of healthy livestock (Folkhälsomyndigheten 2014). We observed a corresponding decrease in total AMR load in the microbiome of brown bears since the 2000s, compared to earlier time periods. Our observation is mirrored by results from screening of healthy farm animals in Sweden for resistance in *Escherichia coli* and *Staphylococcus aureus* isolates, which show low and generally stable or decreasing resistance over the last ten years (Folkhälsomyndigheten and SVA 2019). There is some evidence that restricting antibiotic use in food animals reduces the presence of resistant bacteria in both the animals themselves and human populations in close proximity (Tang et al. 2017). However, how such restrictions affect the surrounding ecosystems and associated wildlife is currently unclear (Goulas et al. 2020).

It is interesting to consider the time lag between the introduction of antibiotic control strategies and an observable reduction in AMR in the environment. It took 10-15 years from the start of antibiotic regulation policies in Sweden in 1985 to see a significant decrease in the AMR levels in the oral microbiome of wild brown bears. The presented pattern should however be interpreted with caution. Wild animal species with different ecology, ranging patterns and affinity to human-modified environments may show important variations of the observed trend. Nevertheless, our study suggests that human actions, both negative and positive, have a direct impact on the environment, including wild animals, and provide hope that the increasing threat from multidrug-resistant bacteria can be turned back following suitable and large-scale policy changes.

### Widespread AMR contamination of natural environments

While total AMR load in brown bears reflected Sweden’s general antibiotic usage, factors related to bear proximity to human activities, such as historical human population density and the 2009 Human Footprint Index, were not associated with AMR load in bear dental calculus. We did not detect any other geographic signal in the data, e.g. related to sample collection coordinates or collection region (municipality/county). This may appear surprising in the face of many studies reporting an association between environmental AMR abundance and proximity to humans or human settlements. For example, a study of AMR in Antarctic seawater found increased AMR levels at sites closer to human habitation (Miller et al. 2009). Multiple studies have shown that wild rodents residing close to livestock have higher AMR levels compared to wild rodents in natural areas (Kozak et al. 2009; Guenther et al. 2010; Grall et al. 2015; Nhung et al. 2015). Associations between AMR and proximity to humans have also been found in large wild mammals, such as foxes, wild boars, deer and tapirs (Skurnik et al. 2006; Cristóbal-Azkarate et al. 2014; Mo et al. 2018). However, our brown bear specimens were generally collected from regions with low historical human population densities (mean bears home range = 4.1 people per km^2^ versus average across Sweden in year 2000 = 21.6 people per km^2^; see Methods) and low Human Footprint indices (Supplementary Figure S9). We therefore may not have observed a strong impact of human proximity on AMR levels because our samples were collected from very similar, little-affected regions.

There is some evidence that AMR contamination can be more widespread than the immediate proximity of source sites. AMR from a likely anthropogenic source has been detected in pristine environments that are expected to be free from human activities, for example, in High Arctic soils and wild birds (Sjölund et al. 2008; McCann et al. 2019). Migratory birds have been suggested as one means for the dissemination of human-associated AMR to remote locations (Arnold et al. 2016). Water and wind-blown soil particles are also likely important substrates for AMR transmission, possibly allowing for dissemination across wide distances (Allen et al. 2010; Taylor et al. 2011). For example, river water and estuarine sediments have been shown to contain AMR, particularly from locations downstream of wastewater treatment plants, livestock farms and other anthropogenic constructions (Mariano et al. 2009; Pruden et al. 2012; He et al. 2016; Zhu et al. 2017; Khan et al. 2019). Furthermore, AMR has been shown to persist in environments even in the absence of selective pressure from antibiotics. Fitness costs for maintaining AMR can be extremely variable and the strength of selection for AMR can remain constant across a large range of antibiotic concentrations, which may facilitate the maintenance of AMR in environmental reservoirs (Vogwill and Maclean 2015; Murray et al. 2018). Our results in brown bears support the idea of widespread environmental contamination with human-made antibiotics and/or resistant bacteria. Presence of antibiotics in the environment will result in increased AMR selective pressures on host-associated microbial communities in wildlife species, while resistant bacteria could be directly integrated into these communities.

### Wildlife as a reservoir for AMR

In this study we focused on host-associated microorganisms, thus targeting ARGs in bacteria that can reside in a mammalian host and are likely more relevant when considering wildlife reservoirs of AMR of potential importance to humans. Resistant bacteria can be transmitted through direct or indirect contact between humans, wildlife and domestic animals via a number of vectors. For example, flies have been identified as a possible vector for transmission of resistant bacteria between humans, domestic animals and environmental samples in both equestrian centres and pig farms (Literak et al. 2009; Dolejska et al. 2011). Wild birds have also been identified as a mechanism of transferring resistant bacteria to livestock, including by physical movement of contaminated material (Luque et al. 2009; Carlson et al. 2015). Less evidence exists for potential exchange of AMR between mammalian wildlife, livestock and humans. However, in Tanzania, AMR prevalence was found to be similarly distributed across humans, domestic animals, wildlife and the environment, likely driven by general transmission of bacteria (Subbiah et al. 2020). In Europe, similar AMR profiles and higher levels of pathogenic bacteria have been found in wild mammals co-habiting with cattle (Leatherbarrow et al. 2007; Navarro-Gonzalez et al. 2012). It is thus becoming clear that wildlife and its role as a potential AMR reservoir should be considered in the global dynamics of AMR and multidrug resistant bacteria.

Omnivorous species have been shown to harbor greater loads of resistant bacteria, likely as a result of their dietary habits and general greater proximity to human settlements (reviewed in (Vittecoq et al. 2016)). Consistently, wild boar, red fox and hedgehog appear to be reservoirs for AMR (Literak et al. 2010; Botti et al. 2013; Radhouani et al. 2013; Zottola et al. 2013; Dias et al. 2015; Bengtsson et al. 2017; Mo et al. 2018). In this study, we focused on another omnivore, the Scandinavian brown bear, as a proxy for wildlife exposure to human-made antibiotics. However, in contrast to the above species, bears are solitary with little direct interaction with humans or livestock. Historically, the Swedish bear population has been small, as it was hunted almost to extinction in the 1930s, before the population increased again following implementation of conservation measures (Swenson et al. 1995). Thus, the historical and modern Swedish brown bears represent a low-density wildlife population without a particular affinity to human settlements. We therefore interpret the temporal changes of AMR levels in the brown bear samples as most likely reflecting the changing widespread contamination of the environment with ARGs capable of persisting in mammalian hosts.

### Dental calculus as a tool for historical AMR research

Using dental calculus from museum specimens, we were able to ‘travel back in time’ to the pre-antibiotics era and quantify temporal changes in AMR load and ARG diversity. As a calcified microbial biofilm, dental calculus is superbly suited for studies of metagenomes from the past. Only few other host-associated microbiomes persist through time, such as coprolites – fossilised faeces that preserve a component of the gut microbiome. However, compared to dental calculus, coprolites are more prone to post-mortem bacterial contamination, have lower DNA preservation and are less abundant in museum and archaeological collections (Warinner et al. 2015).

Despite the great potential of historical microbiomes, one major limitation of metagenomic analyses of historical dental calculus is the possibility of modern contamination. We followed rigorous ancient DNA procedures and limited our study of AMR to oral bacteria, to reduce the possibility of including modern contaminants. However, our classification of ‘oral bacteria’ was primarily based on knowledge of the oral microbial community of humans and pets (Chen et al. 2010; Dewhirst et al. 2012; Dewhirst et al. 2015; Mann et al. 2018) and thus excluded any bacteria which could be specific to the oral cavity of bears (i.e. not carried by humans). As with any metagenomic study from non-model organisms that relies on taxonomic classification using reference databases, a detailed characterisation of the oral microbial community of living brown bears would be a valuable resource and greatly improve our inferences. The degraded nature and low quantity of DNA recovered from historical samples like dental calculus can potentially impact ARG identification. However, we accounted for both DNA fragment length and sequencing depth in our analyses and neither were found to drive the temporal patterns in AMR levels.

One important next step in the study of AMR from dental calculus is the functional validation and characterisation of the detected ARGs. Isolation of bacteria in culture is not possible from historical dental calculus, as the biofilm undergoes periodic mineralisation. However, functional characterisation of detected ARGs could be possible with screening methods that have been successfully applied in both modern and ancient metagenomic studies of AMR (Dcosta et al. 2011; Tsukayama et al. 2018). Historical wildlife microbiomes could also represent an unexplored source of biologically active substances, including novel antibiotics, that could be identified from metagenomes using a strategy targeting biosynthetic gene clusters (Sugimoto et al. 2019). Taken together, the field of ancient metagenomics can broaden our understanding of global environmental trends, including those resulting from human actions, help us evaluate the effectiveness of environmental policies and provide a means to uncover potentially novel active substances that can be recovered from ancient microbial communities.

## Materials and Methods

### Specimens and sample collection

Dental calculus was collected from Swedish brown bear (*Ursus arctos*) specimens from the Swedish Natural History Museum (Stockholm, Sweden). Skulls were macroscopically examined for dental calculus deposits and evidence of oral diseases, such as caries, inflammation and tooth loss. Calculus was removed from the surfaces of teeth without macroscopic signs of oral disease (n=82) with disposable sterile scalpel blades and deposited in sterile microcentrifuge tubes. For most individuals, calculus deposits from multiple teeth were pooled. However, 3 individuals were sampled twice and processed as separate samples, of which 5 samples were excluded due to low oral microbial content. To monitor museum environmental contamination, we used a sterilised cosmetic swab to rub an interior corner of one bear specimen drawer for 5 seconds, before depositing the swab tip in a sterile microcentrifuge tube. To characterise microbial communities that colonise the external surface of museum specimens but are not oral in origin, we repeated the procedure with a new swab on a bear specimen skull, swabbing the cranial surface away from the jaw to avoid potential carryover of endogenous oral microorganisms.

Specimen collection year and location information was obtained from museum records (Supplementary Table S7). For three specimens, year of collection was unknown, however we were able to assign them to the ‘pre 1951’ category based on museum records and information about the collector who donated the specimen. Specimen location information included county (or historical province), municipality, locality description and coordinates. Coordinates were available for the majority of specimens collected after 1995 and were provided in either the RUBIN (RUtin för Biolologiska INventeringar) or Swedish Grid (RT-90) system. They were converted to standard World Geodetic System 84 (WGS84) coordinates using an online converter (http://ormbunkar.se/koordinater/, accessed 10-10-2019). Specimens without location coordinates were assigned WGS84 coordinates by searching for the locality described in the museum record on Google Maps (https://www.google.com/maps, accessed 10-10-2019). Due to changing administrative boundaries over time, we then converted museum records of county/province and municipality to the modern county and municipality based on sample coordinates. Seven specimens had unknown localities (beyond their county of origin) and were excluded from the geography and human impact analyses. However, for visualisation (Figure 1 and Supplementary Figure S10), they were given arbitrary coordinates within their known county.

### Sample processing and DNA extraction

All laboratory protocols were performed in a dedicated ancient DNA laboratory following stringent procedures to minimise contamination (Key et al. 2017). Samples were randomised to control for batch effects and extracted in batches of 16, including two blank negative controls per batch that were taken forward to library preparation. Surface decontamination of dental calculus samples, ranging in weight from < 5 mg and up to 20 mg, consisted of UV light exposure (10 min at 254 nm) followed by a wash in 500 µl of 0.5M ethylenediaminetetraacetate (EDTA) for 1 minute (Brealey et al. 2020). After centrifugation at 18,000 x g for 1 min, the pellet was taken forward for DNA extraction following a silica-based method (Dabney et al. 2013) that was previously successfully applied to non-human dental calculus (Brealey et al. 2020). We eluted purified DNA in 45 µl of EB buffer (10 mM tris-hydrochloride (pH 8.0) (Qiagen, The Netherlands) supplemented with 0.05% (v/v) Tween-20.

### Library preparation and sequencing

We used a double-indexing double-barcoding approach (Rohland et al. 2015; van der Valk et al. 2019) during double-stranded Illumina library preparation (Meyer and Kircher 2010) to guard against index hopping and to retain certainty about sample of origin. We ligated adapters containing inline 7 bp barcodes (Supplementary Table S7) to both ends of the blunt-ended DNA and quantified the incomplete libraries using a real-time PCR assay (primer sequences in Supplementary Table S8) to estimate the number of indexing PCR cycles needed for sequencing (Supplementary Table S7). All extraction and library blanks were consistently lower in DNA content than the majority of samples, as measured by real-time PCR (Supplementary Table S7). Samples with adapter-ligated library concentrations similar to the blanks were excluded. Libraries were double-indexed with unique P5 and P7 indices so that each sample had a unique barcode-index combination (Supplementary Table S7). The first batch of indexing PCR reactions (22 samples and blanks) was performed with 18 µl of adapter-ligated library, 1 µl PfuTurbo C_x_ hotstart polymerase (2.5 U/µl, Agilent Technologies, CA), 5 µl 10X PfuTurbo C_x_ reaction buffer, 0.5 µl dNTP mix (25 mM) and 1 µl of each indexing primer (10 µM) in 50 µl reactions. Following optimisation, the remaining indexing batches were performed with 3.2 µl 10X PfuTurbo C_x_ reaction buffer and the addition of 0.09 µl 20 mg/ml BSA (Thermo Fisher Scientific, CA). For all batches, the PCR cycling conditions were: 2 min at 95^°^C, 8 or 10 cycles (primer sequences in Supplementary Table S8) of 30 sec at 95^°^C, 30 sec at 59^°^C and 1 min at 72^°^C, and a final step of 10 min at 72^°^C. Following purification with MinElute, the indexed libraries were quantified using a real-time PCR assay (primer sequences in Supplementary Table S8). We pooled 1.5 µl of each indexed library (including blanks and swabs) and performed size selection for fragments approximately 100-500 bp in length with AMPure XP beads (Beckman Coulter, IN). The final pooled library was quantified with a Qubit High Sensitivity fluorometer and the fragment length distribution evaluated with the 2200 TapeStation system. The pooled library was sequenced by SciLifeLab Uppsala on 2 Illumina NovaSeq S2 flowcells using paired-end 100 bp read length v1 sequencing chemistry together with additional samples that were not part of this project.

### Data processing

Sequenced reads were demultiplexed and assigned to each sample with an in-house python script based on the unique combination of barcodes and indices, discarding reads with wrong barcode combinations that could be the result of index hopping (van der Valk et al. 2019). Paired-end reads that overlapped by at least 11 bp were merged, adapters and low quality terminal bases (phred scores ≤ 30) were removed and the trimmed, merged reads were filtered to remove reads with a length < 30 bp with AdapterRemoval v2.2.2 (Schubert et al. 2016). Barcode sequences were removed from the 5’ and 3’ ends of merged reads with an in-house python script. All reads for each sample (i.e. across the four lanes from the two Illumina NovaSeq flowcells) were concatenated into a single file per sample. Reads with mean base quality < 30 were filtered out with PrinSeq-Lite v0.20.4 (Schmieder and Edwards 2011). Duplicate reads were removed by randomly keeping one read among those reads having an identical sequence using an in-house python script. To remove any erroneous reads from the Illumina sequencing control phage PhiX, reads were mapped to PhiX (accession: GCA_000819615.1) with bwa mem v0.7.17 (Li and Durbin 2009; Li 2013) and the unmapped reads retained with SAMTools v1.9 (Li et al. 2009) and BEDTools v2.27.1 (Quinlan and Hall 2010). To remove reads originating from the host organism and from human contamination, we mapped reads to a combined reference consisting of the human genome (Schneider et al. 2017) (RefSeq accession: GCF_000001405.38) and the grizzly bear genome (Taylor et al. 2018) (*U. arctos horribilis*, GCF_003584765.1) with bwa mem. The unmapped reads were retained with SAMTools and BEDTools for downstream analyses.

### Microbial source identification

The unmapped reads were assigned taxonomy using the *k*-mer based classifier Kraken2 v2.0.8 (Wood and Salzberg 2014) with the standard Kraken2 database (all archaea, bacteria, viruses and the human genome in RefSeq; built 2019-05-01) and default parameters. We used Kraken-biom (github.com/smdabdoub/kraken-biom) to extract the summarised number of reads assigned at the genus and species levels. These assignments were used with SourceTracker v1.0 (Knights et al. 2011) in R, to estimate the potential contribution of source microbiomes to our samples. Source sequencing reads were processed through the same pipeline as sample reads, and included soil (Johnston et al. 2016), human skin (Oh et al. 2014), human gut (Huttenhower et al. 2012; Lloyd-price et al. 2018), human supragingival plaque (Huttenhower et al. 2012; Lloyd-price et al. 2018), human medieval dental calculus (Mann et al. 2018) and laboratory reagent (Salter et al. 2014) microbiomes (Supplementary Table S9). Within SourceTracker, sample rarefaction was set to the source with the lowest summed sequencing depth (-r 20809).

### Oral bacteria identification

We used Bracken v2.0 (Lu et al. 2017) to estimate taxa abundances from the Kraken read assignments at the species level (-l S) using a read length of 65 bp (-r 65), a k-mer length of 35 bp (-k 35) and without an abundance threshold (-t 0). The data were further processed in RStudio v1.3.959 (RStudio Team 2020) using R v4.0.2 (R Core Team 2020). We excluded bear calculus samples with low microbial taxa abundances (summed Bracken abundance < 12,390, corresponding to the summed Bracken abundance of the most deeply sequenced extraction blank). To reduce false-positive taxonomic assignments, we filtered out taxa present at < 0.05% relative abundance (Bracken abundance divided by sum of Bracken abundance in a sample) (Velsko et al. 2018). We also excluded calculus samples with low proportions of taxa showing similarity to human oral microbiomes (< 5% of a sample attributed to human calculus and plaque), as estimated by SourceTracker, to remove potentially strongly contaminated samples. Since we were interested in endogenous AMR of the host-associated oral microbiome and environmental microorganisms may contain ARGs, we subset the dataset to a list of oral bacteria (Supplementary Table S2). To this end, we used previously defined criteria (Brealey et al. 2020) and classifications from (Mann et al. 2018), taxa present in the Human Oral Microbiome Database (Chen et al. 2010) and those identified in dog and cat oral microbiota studies (Dewhirst et al. 2012; Dewhirst et al. 2015). Bracken relative abundances of oral taxa were then summarised to the genus level for some downstream analyses.

### Antimicrobial resistance profiling

AMR profiling was performed as previously described (Brealey et al. 2020). Oral taxa reads were blasted against the Comprehensive Antibiotic Resistance Database (CARD) v3.0.1 (modified 2019-02-19) (Jia et al. 2017), a curated collection of resistance determinant sequences, with blast v2.0.9+ (Altschul et al. 1990; Madden et al. 2009) using default parameters. Using RStudio, the DNA and Protein accession numbers associated with each CARD sequence were mapped to their respective Antibiotic Resistance Ontology (ARO) accession number and used to obtain the AMR gene family (ARG family) and resistance mechanism of each sequence. The best hit for a read was identified based on highest bit score. Where multiple hits had the same bit score, we compared the ARO terms and in all cases, the hits shared the same ARO information (ARG family and resistance mechanism). We therefore randomly chose one hit to carry forward. Total AMR load was calculated as the sum of all reads with hits in a sample, normalised by the number of sequenced oral bacteria reads. ARG family abundance was calculated as the sum of reads assigned to each ARG family in a sample normalised by the number of sequenced oral bacteria reads in that sample. We used the R package vegan (Oksanen et al. 2018), function ‘specnumber’, to calculate ARG diversity per sample as the number of ARG families in a sample with at least one sequencing read. To obtain ARG family abundance across a time period, we calculated the sum of reads with a best hit for each ARG family in a time period (i.e. combined across samples) normalised by the number of extracted oral reads in all samples in that time period. Similarly, ARG diversity per time period was calculated as the number of ARG families in a time period with at least one sequencing read (i.e. combined across samples).

### Geographic and human impact data

Historical human population data was based on publicly available Swedish church parish (församling or socken in Swedish) records. Parishes were both religious and territorial units until 1995, and recorded parish population numbers, among other details. The 50 specimens with known localities were plotted in QGIS v3.14.15 (QGIS.org 2020) with a vector of Swedish parish boundaries as they were in the 1976-1995 Swedish property register, downloaded from Lantmäteriet, the Swedish mapping, cadastral and land registration authority (ftp://download-opendata.lantmateriet.se/SockenStad/Sverige/Sweref_99_TM/Geopackage/snst_riks_Sweref_99_TM_gpkg.zip, accessed 27-05-2020). The area of each parish was calculated using the ‘Add Geometry Attributes’ tool in QGIS. The corresponding parish for each sample was determined using the tool ‘Join Attributes by Location’ (Geometric predicate: ‘intersects’, ‘within’; Join type: take attributes of the feature with the largest overlap (one-to-one)). Publicly available historical human population data was downloaded using the FOLKNET search tool (http://rystad.ddb.umu.se:8080/FolkNet/index.jsp, accessed 08-06-2020). The FOLKNET database contains the human population from each town and parish, collected every 10 years from 1810 to 1990.

Secular municipality divisions came into effect between 1971 and 1995 and the Swedish Tax Agency took over the population register in 1991. Therefore, modern human population data (for samples collected between 1995 and 2016) was based on publicly available population statistics of Swedish towns from Statistics Sweden (SCB). ArcView GIS shape files for human population in 2000, 2010 and 2015 were downloaded (https://www.scb.se/vara-tjanster/oppna-data/oppna-geodata/tatorter/ and https://www.scb.se/vara-tjanster/oppna-data/oppna-geodata/smaorter/, accessed 09-06-2020). In QGIS, the two shape file vectors were merged for each reference year (‘Merge vector layers’) and cleaned-up (‘Check validity & Fix geometries’). Each merged vector contained the polygon boundaries and area of each town with its human population on 31st of December of the reference year. Because historical parish boundaries tended to be larger than modern town boundaries, we chose to map the modern towns to the historical parish boundaries using the ‘Join Attributes by Location’ tool. Thus, data from multiple towns within a parish boundary were combined. The historical parishes and corresponding modern town names, populations and areas were exported into RStudio and summed across historical parish for each reference year.

In RStudio, specimens were assigned to their closest reference year (available for each decade, e.g. a specimen collected in 1984 was assigned to the reference year 1980, while a specimen collected in 1985 was assigned to 1990), except for samples collected after 2014, which were assigned to the 2015 reference year. For samples collected before 1995, human population was assigned as the FOLKNET historical population for the parish during the closest reference year, whereas samples collected from 1995 were assigned human population values based on the information from SCB. Human population density (population per square kilometre) was then calculated using the area of each parish. For Supplementary Figure S9, human population density for all of Sweden and for each county in the year 2000 was downloaded from SCB (https://www.statistikdatabasen.scb.se/pxweb/en/ssd/STARTBEBE0101BE0101C/BefArealTathetKon/, accessed 14-12-2020).

Human activities data were not consistently available for historical time periods. We therefore used the Last of the Wild Project (version 3) 2009 Human Footprint, 2018 Release, which is a global map of the cumulative human pressure on the terrestrial environment in 2009, using eight variables (built-up environments, population density, electric power infrastructure, crop lands, pasture lands, roads, railways and navigable waterways) at a spatial resolution of approximately 1 km (Venter et al. 2016). The GeoTiff raster file for 2009 was downloaded from the NASA Socioeconomic Data and Applications Center (https://doi.org/10.7927/H46T0JQ4, accessed 04-06-2020) and imported into QGIS with the 50 specimens with assigned coordinates (visualised in Figure 1). A 500 km^2^ buffer zone, corresponding to a 12.5 km radius, was calculated around each sample, corresponding to an approximate average of the home range size of adult brown bears in Sweden (100-1000 km^2^, depending on sex and reproductive status) (Dahle and Swenson 2003). The average Human Footprint Index was calculated within each sample buffer zone (the ‘Zonal statistics’ tool) and exported into RStudio. For Supplementary Figure S9, the average Human Footprint Index was also calculated across all of Sweden.

### Statistical analysis

All statistical analyses were performed in RStudio. Specimen time period was treated as a five-level ordered factor for all relevant analysis. For ordination, Bracken abundance counts for all filtered taxa were normalised by the centre-log ratio (CLR) transformation, using a pseudocount of 1 added to all taxa in all samples to resolve the problem of zero values. Euclidean distance matrices were calculated with the vegan (Oksanen et al. 2018) function vegdist. Non-metric multidimensional scaling (NMDS) was performed on the distance matrices with the vegan function metaMDS, with k = 3 for all taxa (Supplementary Figure S2) and k = 2 for oral bacteria (Supplementary Figure S8). NMDS was also performed for oral bacteria on a Jaccard distance matrix calculated from binary presence/absence data with k = 2 (Supplementary Figure S8). PERMANOVA was performed on distance matrices using the vegan function adonis with 1000 permutations (Supplementary Table S1). To evaluate changes in abundance (total AMR load and ARG family abundance) over the time periods, generalised linear models (GMLs) were built with a quasibinomial distribution and a logit link function and total oral read count as weights. To determine which potential confounding factors were associated with total AMR load, continuous explanatory variables (e.g. median length of oral reads) were evaluated with a Spearman correlation test, while categorical variables (e.g. extraction batch) were evaluated with a Kruskal rank sum test (Supplementary Table S3). Significant variables were included as confounding factors in the generalised linear models.

ARG family diversity was calculated across samples pooled by time period, as the number of unique ARG families detected in that time period, using the vegan function specnumber. To control for different numbers of samples between time periods, we subsampled each time period to 8 samples (the lowest number of samples in any time period) without replacement, using the R base function sample, before calculating ARG family diversity. This process was repeated 1000 times to generate a distribution of ARG family diversity values for each time period (Figure 3a). To control for differences in sequencing depth between time periods, we also subsampled each time period to 500 AMR-positive reads (the lowest number of pooled AMR-positive reads in a time period was 553), independent of sample ID, and calculated ARG family diversity 1000 times, as above (Supplementary Figure S5).

To identify correlations between ARG family abundance and bacterial genus abundance, we used the R package CCREPE (Compositionality Corrected by REnormalizaion and PErmutation, https://github.com/biobakery/ccrepe), which aims to identify significant correlations between two compositional datasets while accounting for multiple hypothesis testing. We compared the four most highly abundant ARG families (ABC-F, ileS, RND and rpoB) with all oral bacterial genera, 20 bootstrap and permutation iterations and a minimum non-zero sample (subject) number per ARG family/genus of 20. For the measure of similarity (or correlation) between samples, we used CCREPE’s in-build nc.score function, which calculates species-level co-variation and co-exclusion patterns based on an extension of Diamond’s checkerboard score to ordinal data. Significantly correlated ARG–genus pairs were identified as those with a q value (p value adjusted for multiple hypotheses with the Benjamin-Hochberg-Yekutieli procedure) < 0.05 and a positive correlation (i.e. nc.score > 0).

## Data availability

Raw sequencing data can be found at the European Nucleotide Archive under the project accession PRJEB42014 (sample-specific ENA accessions are provided in Supplementary Table S7). Sample metadata is provided in Supplementary Table S7. In-house python and R scripts used in data processing and analysis are available on GitHub (https://github.com/jcbrealey/amr_bears).

## Acknowledgements

We thank Peter Niehoff, Kevin Mulder and Tom van der Valk for their assistance with sampling and Maria Wisselgren from the Centre for Demographic and Aging Research (CEDAR) at Umeå University for advice on accessing and navigating Swedish historical parish records. Sequencing was performed by the SNP&SEQ Technology Platform in Uppsala. The facility is part of the National Genomics Infrastructure Sweden and Science for Life Laboratory. The SNP&SEQ Platform is also supported by the Swedish Research Council and the Knut and Alice Wallenberg Foundation. We also acknowledge the National Bioinformatics Infrastructure for providing computational resources to this project. We obtained geographic and human population data from a number of publicly available resources: Lantmäteriet (the Swedish mapping, cadastral and land registration authority), SCB (Statistics Sweden) and FOLKNET (provided by CEDAR at Umeå University). This work was supported by the Formas grants 2016-00835 and 2019-00275 and the Science for Life Laboratory National Sequencing Projects grant (NP00039) to KG.

## Competing interests

The authors declare no competing interests.

## SUPPLEMENTARY MATERIAL

**Supplementary Figure S1.**
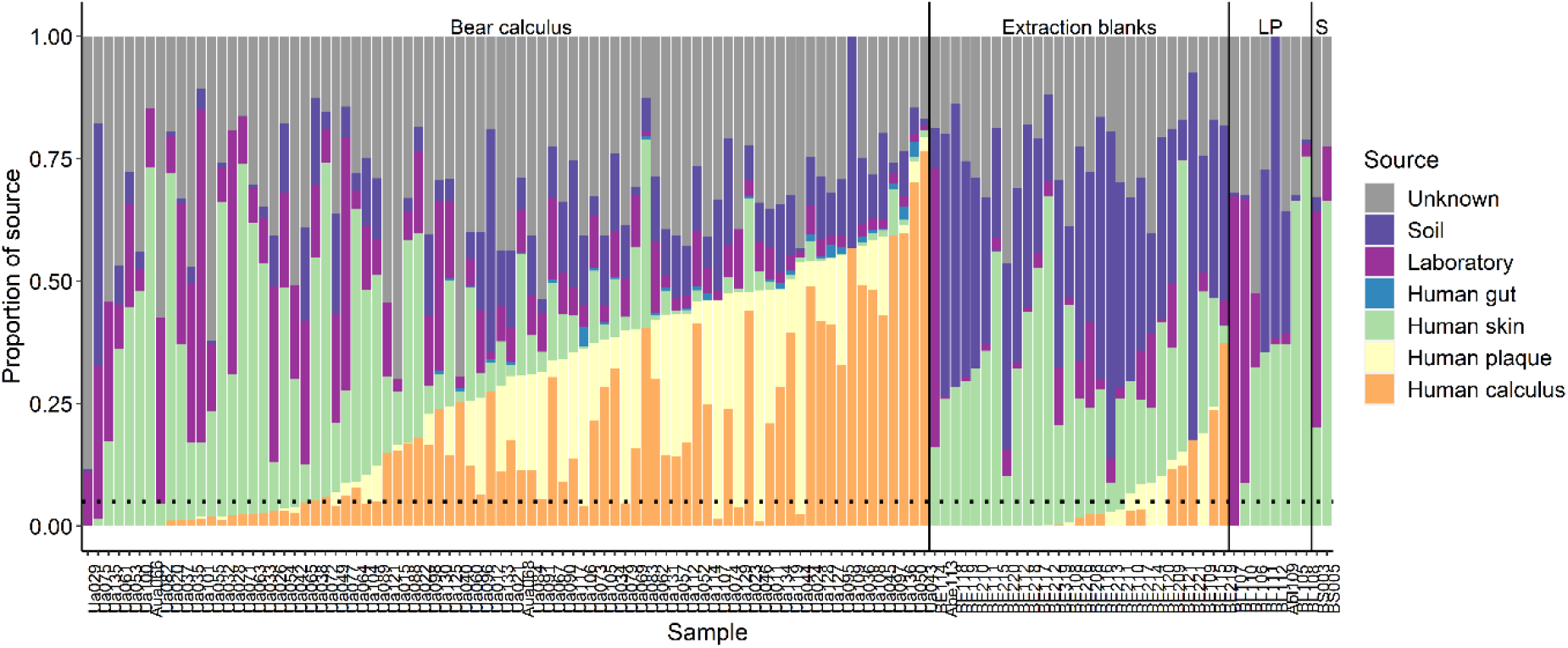
SourceTracker analysis of microbial composition of bear dental calculus samples, extraction and library preparation (LP) blank control samples and museum swabs (S). Microbial taxonomy was assigned by Kraken2. SourceTracker was run prior to additional filtering on samples or taxa. Dental calculus samples with < 5% oral microbial signature (human plaque + human calculus sources) were excluded from further analysis (dashed line).

**Supplementary Figure S2.**
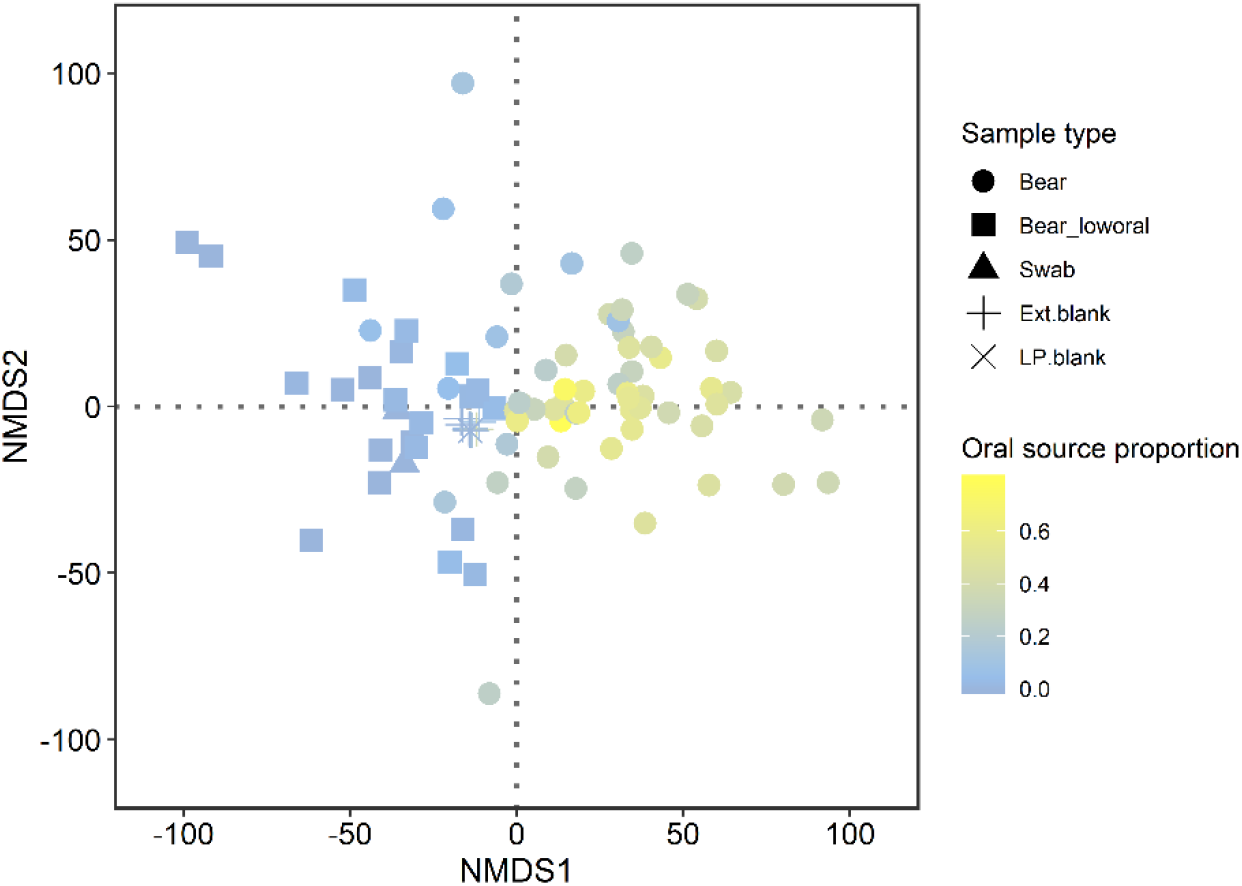
NMDS of CLR-normalised microbial taxa abundances in bear dental calculus, museum swab, extraction blank and library preparation blank samples. NMDS was performed with k = 3 on a Euclidean distance matrix. NMDS stress: 0.111. Samples are coloured by the proportion of their microbiome composition corresponding to human oral microbiome, based on SourceTracker results (summed human dental calculus and human dental plaque proportions). Bear samples that were excluded due to low oral proportions (< 0.05) are indicated by squares instead of circles. In a PERMANOVA of Euclidean distances (Supplementary Table S1), sample type (dental calculus or various controls) accounted for 4.28% of the variation (F_3,112_ = 1.69, p = 0.012) and oral source proportion accounted for 0.83% (F_1,112_ = 0.98, p = 0.405). However, when oral source proportion was included in the PERMANOVA without sample type, it accounted for 1.88% of the variation (F_1,116_ = 2.23, p = 0.021).

**Supplementary Figure S3.**
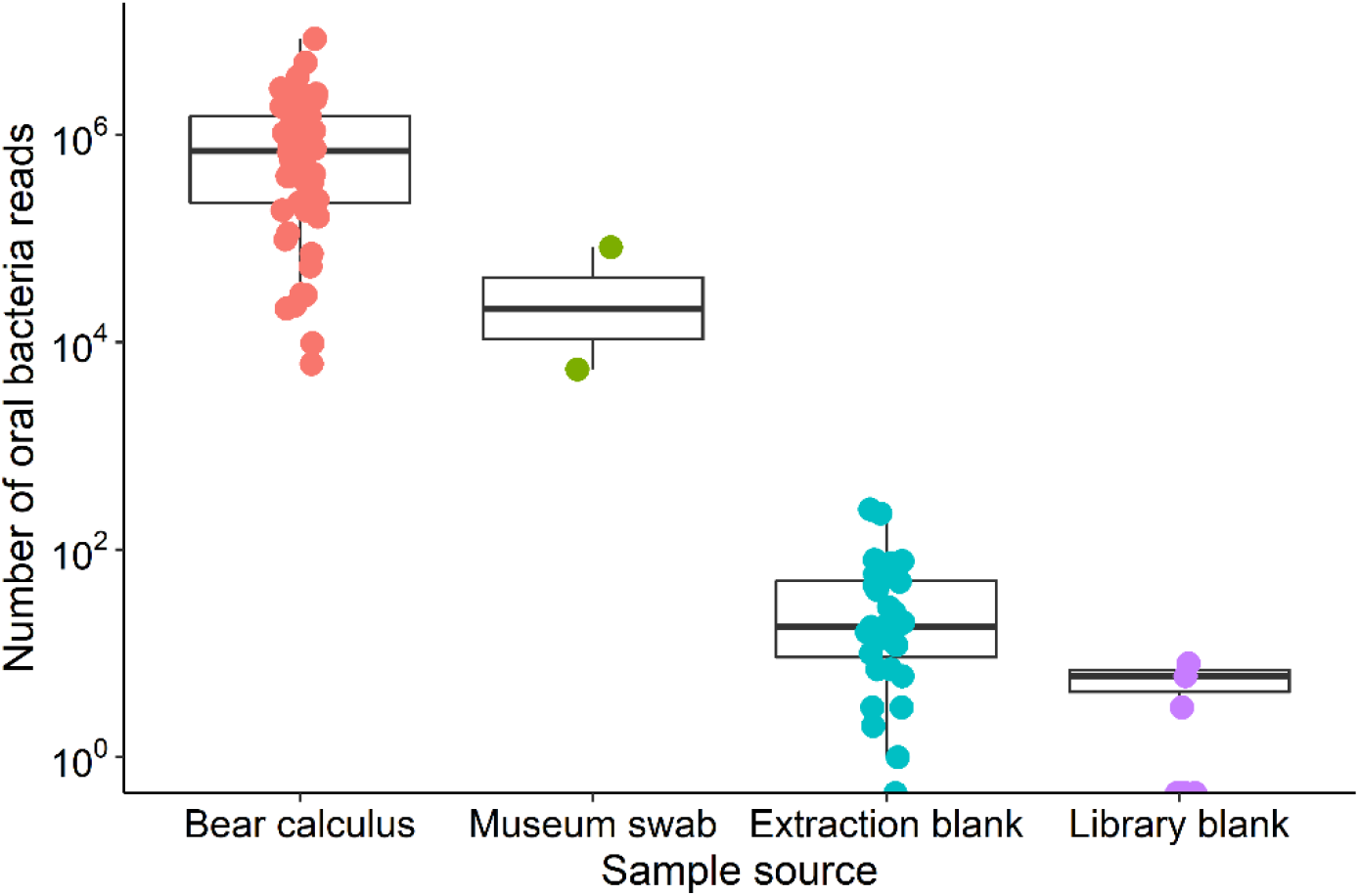
Number of reads assigned to oral bacteria taxonomy (Supplementary Table S2) identified in bear dental calculus samples, museum swabs and laboratory blanks. Number of reads is displayed on a log scale (y-axis).

**Supplementary Figure S4.**
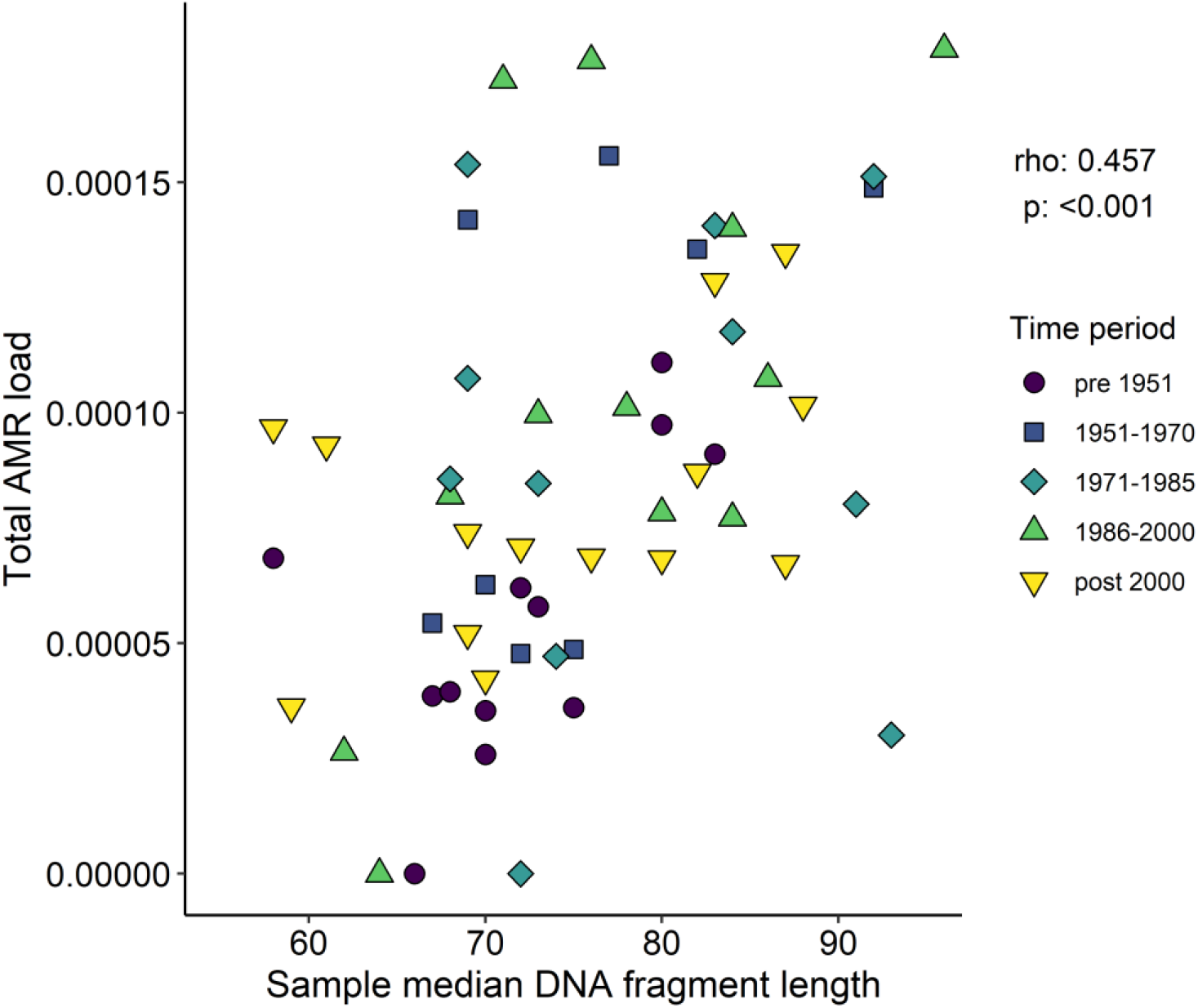
Total AMR load is correlated with median length of oral bacteria DNA fragments in each bear calculus sample. Sample colour and shape indicates sample collection time period. Spearman correlation rho and p are shown above the legend.

**Supplementary Figure S5.**
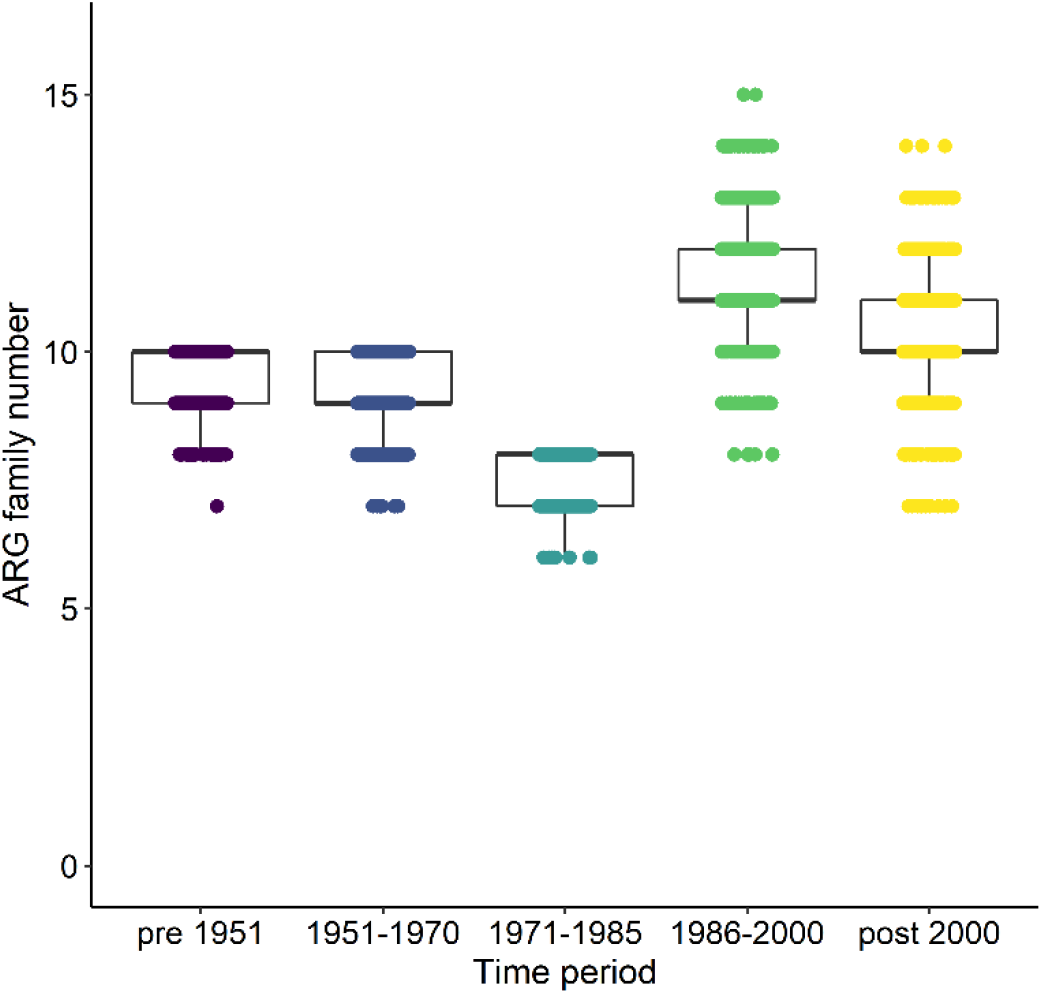
Boxplots of the number of unique ARG families detected in each time period while controlling for differences in the amount of data available between time periods. All ARG-positive reads from all samples in a time period were pooled, subsampled to 500 reads and the number of unique ARG families calculated. This subsampling was independently repeated 1000 times to generate the boxplots.

**Supplementary Figure S6.**
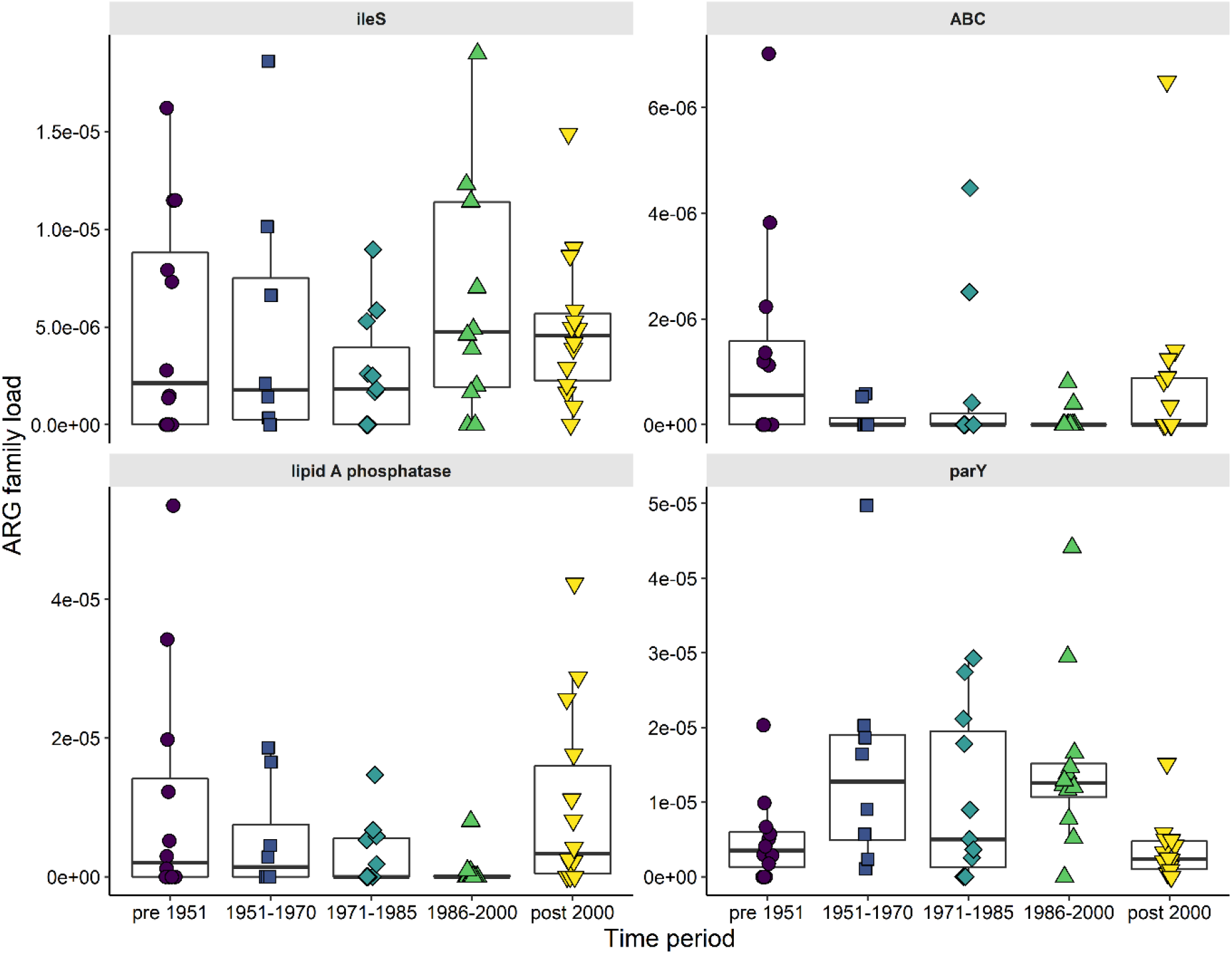
Proportion of oral bacteria reads mapping to four out of seven most abundant ARG families in bear calculus samples across the study time periods (for the remaining three see Figure 4). (a) antibiotic resistant ileS (isoleucyl-tRNA synthestase), (b) ABC (ATP-binding cassette) antibiotic efflux pump, (c) lipid A phosphatase and (d) aminocoumarin resistant parY (topoisomerase IV).

**Supplementary Figure S7.**
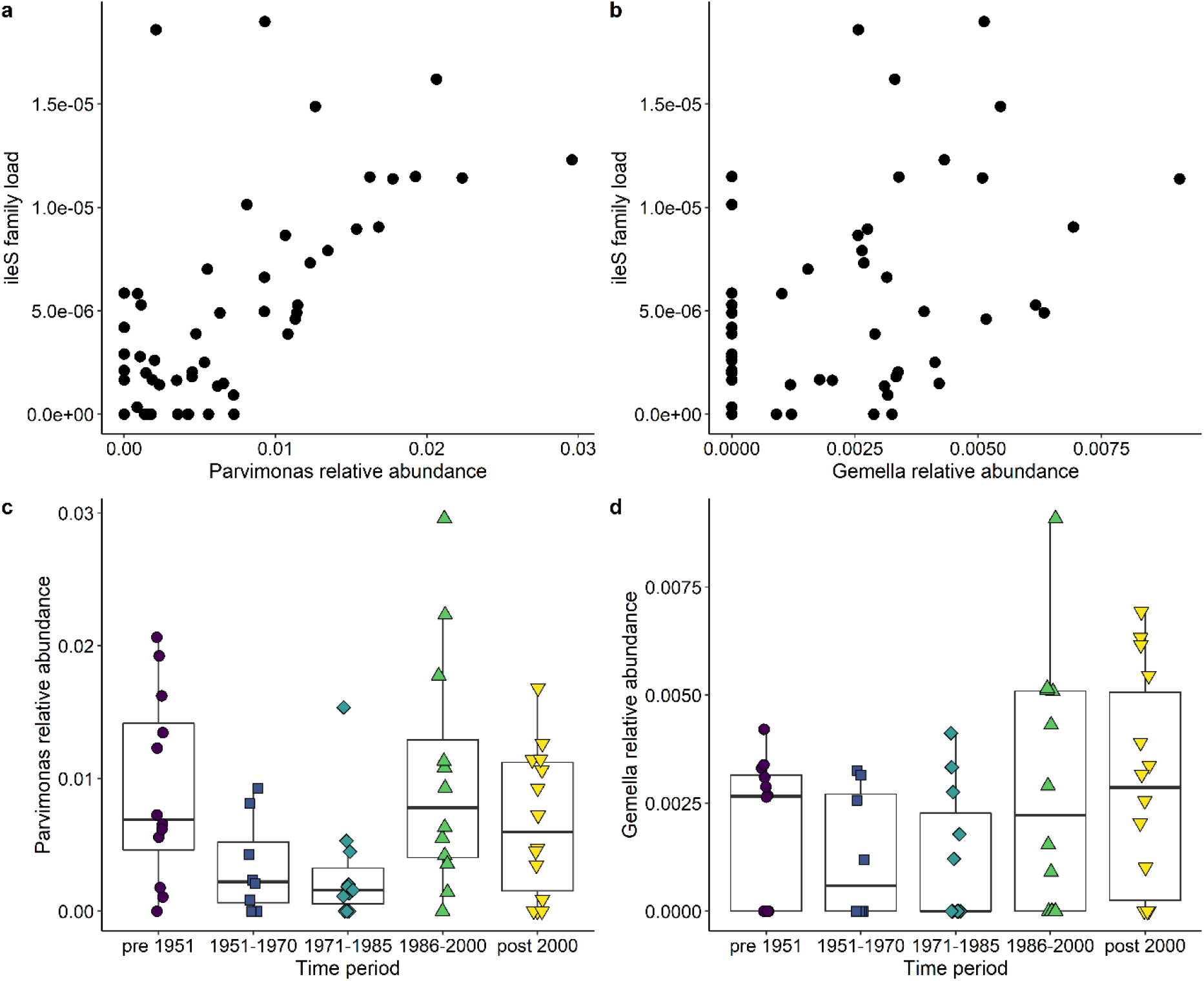
Changes in abundance of antibiotic resistant ileS (isoleucyl-tRNA synthestase) is driven by changes in abundance of *Parvimonas* (a,c) and *Gemella* (b,d). Abundance of ileS is correlated with *Parvimonas* (a) and *Gemella* (b) abundance. Changes in abundance of the bacterial genera *Parvimonas* (c) and *Gemella* (d) over time. ileS abundance was calculated as the proportion of oral bacteria reads mapping to the ileS family. Genus abundance was calculated as the proportion of oral bacteria reads assigned by Bracken to that genus.

**Supplementary Figure S8.**
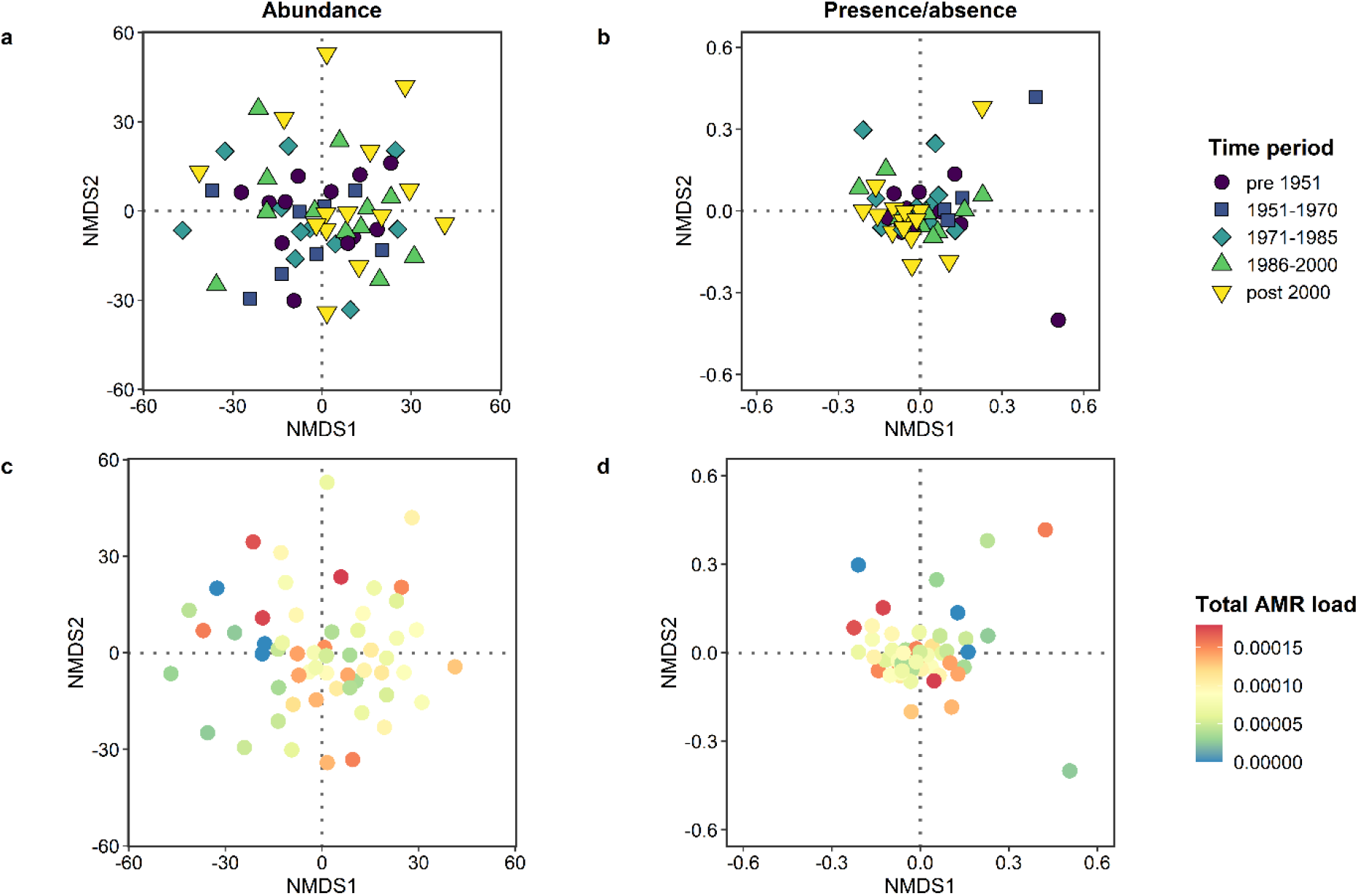
NMDS of CLR-normalised abundances (a,c) and presence/absence (b,d) of oral bacteria in bear dental calculus samples. NMDS was performed with k = 2 on a Euclidean (a,c) or Jaccard (b, d) distance matrix. NMDS stress for (a,b): 0.188 and for (b,d): 0.132. Samples are coloured by sample collection time period (a-b) and total AMR load (c-d). Neither variable accounted for a significant percentage of variation in a PERMANOVA of either distance matrix (Supplementary Table S1).

**Supplementary Figure S9.**
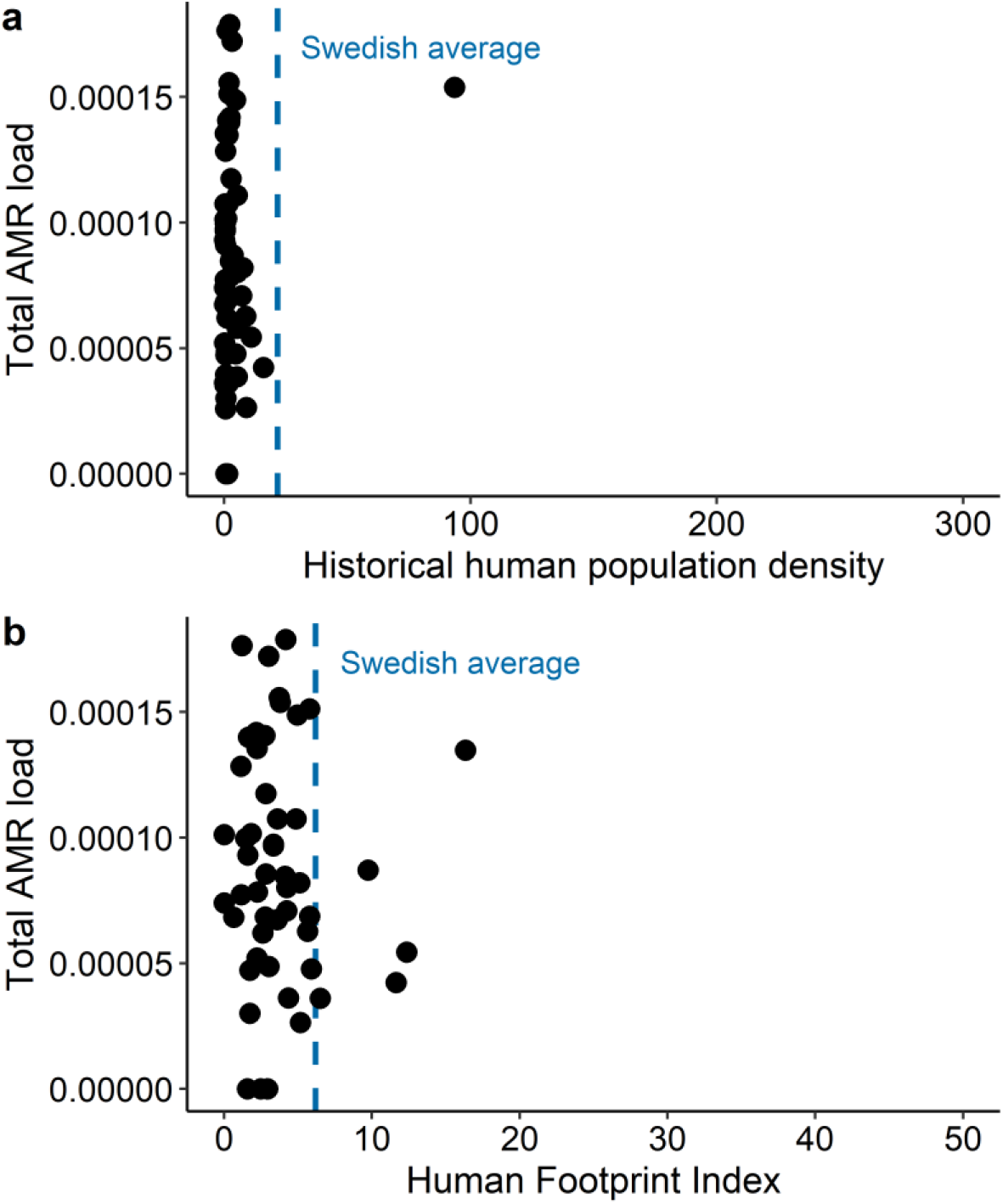
Total AMR load compared to historical human population per km^2^ (a) and mean Human Footprint Index (b) within a 12.5 km radius (500 km^2^) around specimen collection location. The majority of bear specimens were collected from low-density areas, compared to the population density of Sweden as a whole in 2000 (dashed blue line in a) and from areas with low Human Footprint Indices compared to the average across Sweden (dashed blue line in b). The upper limit of the x axis in a is indicative of the most densely populated region in Sweden (Stockholm county, 280 people per km^2^ in 2000) and in b is indicative of the maximum Human Footprint Index in Sweden (50).

**Supplementary Figure S10.**
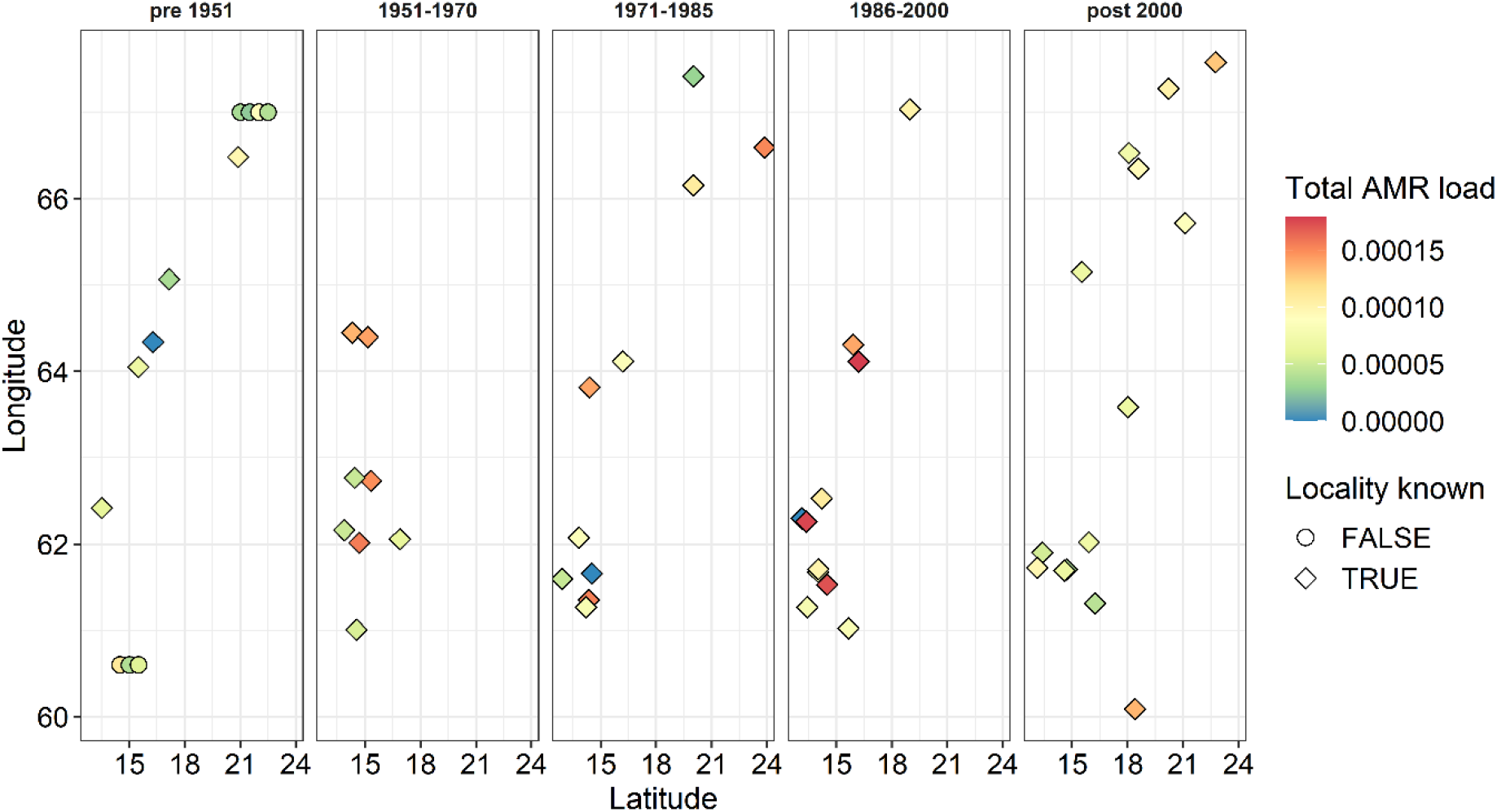
Sample locations (longitude and latitude) divided by sample collection time period and coloured by total AMR load. Seven samples from pre-1951 were missing locality information (indicated by circles instead of diamonds) and are plotted using arbitrary coordinates within their respective counties.

### Supplementary Figures

**Supplementary Table** legends (for tables, see Excel workbook)

Supplementary Table S1. PERMANOVA results of microbial community differences between i) all samples (including laboratory controls), using both oral and non-oral microbial CLR-normalised abundances and Euclidean distances; ii-iii) bear samples using oral microbial CLR-normalised abundances and Euclidean distances; and iv-v) bear samples using oral microbial detection (presence/absence) and Jaccard distances. In ii) and iv), an associated with time period was investigated, while in iii) and v), an association with total AMR load (the number of reads mapping to CARD divided by the total number of oral bacterial reads) was investigated.

Supplementary Table S2. Taxonomy and NCBI taxon IDs of bacteria classified as ‘oral’ for the AMR analyses.

Supplementary Table S3. Tests for associations between total AMR load and variables relating to sampling processing (e.g. processing batches, sequencing depth), specimen collection (e.g. time period, location) and specimen proximity to humans (e.g. historical human population density, Human Footprint Index). Categorical variables were assessed with a Kruskal-Wallis rank sum and continuous variables with a Spearman’s correlation test.

Supplementary Table S4. ARG family descriptions (abbreviation, name and resistance mechanism).

Supplementary Table S5. Results from generalised linear models of changes in the abundance of each of the seven most abundance ARG families (ABC-F, rpoB, RND, ileS, ABC, pgpB and parY) load across time period, controlling for median DNA fragment length.

Supplementary Table S6. Results from generalised linear models of the effect of geography (longitude and latitude), historical human population density and 2009 Human Footprint Index on the total AMR load, with and without controlling for time period and median DNA fragment length.

Supplementary Table S7. Sample metadata.

Supplementary Table S8. Sequencing adapter and real-time PCR assay primer sequences. Supplementary Table S9. Samples used as sources for SourceTracker analysis.

## Notes

### Competing Interest Statement

The authors have declared no competing interest.

## References

Adler CJ, Dobney K, Weyrich LS, Kaidonis J, Walker AW, Haak W, Bradshaw CJ a, Townsend G, Sołtysiak A, Alt KW, et al. 2013. Sequencing ancient calcified dental plaque shows changes in oral microbiota with dietary shifts of the Neolithic and Industrial revolutions. Nat Genet. 45:450–455.

Alekshun MN, Levy SB. 2007. Molecular Mechanisms of Antibacterial Multidrug Resistance. Cell. 128:1037–1050.

Allen HK, Donato J, Wang HH, Cloud-Hansen KA, Davies J, Handelsman J. 2010. Call of the wild: Antibiotic resistance genes in natural environments. Nat Rev Microbiol. 8:251–259.

Altschul SF, Gish W, Miller W, Myers EW, Lipman DJ. 1990. Basic local alignment search tool. J Mol Biol. 215:403–410.

Arnold KE, Williams NJ, Bennett M. 2016. “Disperse abroad in the land”: The role of wildlife in the dissemination of antimicrobial resistance. Biol Lett. 12:20160137.

Begemann S, Perkins E, Van Hoyweghen I, Christley R, Watkins F. 2018. How Political Cultures Produce Different Antibiotic Policies in Agriculture: A Historical Comparative Case Study between the United Kingdom and Sweden. Sociol Ruralis. 58:765–785.

Bengtsson B, Persson L, Ekström K, Unnerstad HE, Uhlhorn H, Börjesson S. 2017. High occurrence of mecC-MRSA in wild hedgehogs (Erinaceus europaeus) in Sweden. Vet Microbiol. 207:103–107.

Bhullar K, Waglechner N, Pawlowski A, Koteva K, Banks ED, Johnston MD, Barton HA, Wright GD. 2012. Antibiotic resistance is prevalent in an isolated cave microbiome. PLoS One. 7:e34953.

Botti V, Valérie Navillod F, Domenis L, Orusa R, Pepe E, Robetto S, Guidetti C. 2013. Salmonella spp. and antibiotic-resistant strains in wild mammals and birds in north-western Italy from 2002 to 2010. Vet Ital. 49:195–202.

Brealey JC, Leitão HG, van der Valk T, Xu W, Bougiouri K, Dalén L, Guschanski K. 2020. Dental calculus as a tool to study the evolution of the mammalian oral microbiome. Mol Biol Evol. 37:3003–3022.

Carlson JC, Hyatt DR, Ellis JW, Pipkin DR, Mangan AM, Russell M, Bolte DS, Engeman RM, DeLiberto TJ, Linz GM. 2015. Mechanisms of antimicrobial resistant Salmonella enterica transmission associated with starling-livestock interactions. Vet Microbiol. 179:60–68.

Cassini A, Högberg LD, Plachouras D, Quattrocchi A, Hoxha A, Simonsen GS, Colomb-Cotinat M, Kretzschmar ME, Devleesschauwer B, Cecchini M, et al. 2019. Attributable deaths and disability-adjusted life-years caused by infections with antibiotic-resistant bacteria in the EU and the European Economic Area in 2015:a population-level modelling analysis. Lancet Infect Dis. 19:56–66.

Chen T, Yu WH, Izard J, Baranova O V., Lakshmanan A, Dewhirst FE. 2010. The Human Oral Microbiome Database: a web accessible resource for investigating oral microbe taxonomic and genomic information. Database (Oxford).

Chitsaz M, Booth L, Blyth MT, O’mara ML, Brown MH. 2019. Multidrug resistance in Neisseria gonorrhoeae: Identification of functionally important residues in the MtrD efflux protein. MBio. 10:e02277–19.

Coats SR, To TT, Jain S, Braham PH, Darveau RP. 2009. Porphyromonas gingivalis resistance to polymyxin B is determined by the lipid A 4’-phosphatase, PGN_0524. Int J Oral Sci. 1:126–135.

Cristóbal-Azkarate J, Dunn JC, Day JMW, Amábile-Cuevas CF. 2014. Resistance to antibiotics of clinical relevance in the fecal microbiota of Mexican wildlife. PLoS One. 9:e107719.

Dabney J, Knapp M, Glocke I, Gansauge M-T, Weihmann A, Nickel B, Valdiosera C, Garcia N, Paabo S, Arsuaga J-L, et al. 2013. Complete mitochondrial genome sequence of a Middle Pleistocene cave bear reconstructed from ultrashort DNA fragments. Proc Natl Acad Sci. 110:15758–15763.

Dahle B, Sørensen OJ, Wedul EH, Swenson JE, Sandegren F. 1998. The diet of brown bears Ursus arctos in central Scandinavia: Effect of access to free-ranging domestic sheep Ovis aries. Wildlife Biol. 4:147–158.

Dahle B, Swenson JE. 2003. Home ranges in adult Scandinavian brown bears (Ursus arctos): Effect of mass, sex, reproductive category, population density and habitat type. J Zool. 260:329–335.

Dcosta VM, King CE, Kalan L, Morar M, Sung WWL, Schwarz C, Froese D, Zazula G, Calmels F, Debruyne R, et al. 2011. Antibiotic resistance is ancient. Nature. 477:457–461.

Dent VE. 1982. Identification of oral Neisseria species of animals. J Appl Bacteriol. 52:21–30.

Dewhirst FE, Klein EA, Bennett ML, Croft JM, Harris SJ, Marshall-Jones Z V. 2015. The feline oral microbiome: A provisional 16S rRNA gene based taxonomy with full-length reference sequences. Vet Microbiol. 175:294–303.

Dewhirst FE, Klein EA, Thompson EC, Blanton JM, Chen T, Milella L, Buckley CMF, Davis IJ, Bennett ML, Marshall-Jones Z V. 2012. The canine oral microbiome. PLoS One. 7:1–12.

Dias D, Torres RT, Kronvall G, Fonseca C, Mendo S, Caetano T. 2015. Assessment of antibiotic resistance of Escherichia coli isolates and screening of Salmonella spp. in wild ungulates from Portugal. Res Microbiol. 166:584–593.

Dolejska M, Duskova E, Rybarikova J, Janoszowska D, Roubalova E, Dibdakova K, Maceckova G, Kohoutova L, Literak I, Smola J, et al. 2011. Plasmids carrying blaCTX-M-1 and qnr genes in Escherichia coli isolates from an equine clinic and a horseback riding centre. J Antimicrob Chemother. 66:757–764.

Edqvist LE, Pedersen KB. 2001. Antimicrobials as growth promoters: Resistance to common sense. In: edHarremoës P, Gee D, MacGarvin M, Stirling A, Keys J, Wynne B, Vaz SG, editors. Late lessons from early warnings: the precautionary principle 1896-2000. Copenhagen: European Environment Agency. p. 100–110.

Ekdahl K, Hansson HB, Mölstad S, Söderström M, Walder M, Persson K. 1998. Limiting the spread of penicillin-resistant Streptococcus pneumoniae: Experiences from the south Swedish pneumococcal intervention project. Microb Drug Resist. 4:99–105.

Elfström M, Davey ML, Zedrosser A, Müller M, De Barba M, Støen OG, Miquel C, Taberlet P, Hackländer K, Swenson JE. 2014. Do Scandinavian brown bears approach settlements to obtain high-quality food? Biol Conserv. 178:128–135.

Filippova SN, Surgucheva NA, Kolganova T V., Cherbunina MY, Brushkov A V., Mulyukin AL, Gal’chenko VF. 2019. Isolation and Identification of Bacteria from an Ice Wedge of the Mamontova Gora Glacial Complex (Central Yakutia). Biol Bull. 46:234–241.

Folkhälsomyndigheten. 2014. Swedish work on containment of antibiotic resistance.

Folkhälsomyndigheten, SVA. 2019. Swedres-Svarm 2018. Consumption of antibiotics and occurrence of resistance in Sweden. Solna/Uppsala.

Francino MP. 2016. Antibiotics and the human gut microbiome: Dysbioses and accumulation of resistances. Front Microbiol. 6:1543.

Fusté E, Galisteo GJ, Jover L, Vinuesa T, Villa TG, Viñas M. 2012. Comparison of antibiotic susceptibility of old and current Serratia. Future Microbiol. 7:781–786.

Gibbs EPJ. 2014. The evolution of One Health: a decade of progress and challenges for the future. Vet Rec. 174:85–91.

Goethem MW Van, Pierneef R, Bezuidt OKI, Van De Peer Y, Cowan DA, Makhalanyane TP. 2018. A reservoir of ‘historical’ antibiotic resistance genes in remote pristine Antarctic soils. Microbiome. 6:40.

Goulas A, Belhadi D, Descamps A, Andremont A, Benoit P, Courtois S, Dagot C, Grall N, Makowski D, Nazaret S, et al. 2020. How effective are strategies to control the dissemination of antibiotic resistance in the environment? A systematic review. Environ Evid. 9:4.

Grall N, Barraud O, Wieder I, Hua A, Perrier M, Babosan A, Gaschet M, Clermont O, Denamur E, Catzeflis F, et al. 2015. Lack of dissemination of acquired resistance to β-lactams in small wild mammals around an isolated village in the Amazonian forest. Environ Microbiol Rep. 7:698–708.

Guenther S, Grobbel M, Heidemanns K, Schlegel M, Ulrich RG, Ewers C, Wieler LH. 2010. First insights into antimicrobial resistance among faecal Escherichia coli isolates from small wild mammals in rural areas. Sci Total Environ. 408:3519–3522.

He LY, Ying GG, Liu YS, Su HC, Chen J, Liu SS, Zhao JL. 2016. Discharge of swine wastes risks water quality and food safety: Antibiotics and antibiotic resistance genes from swine sources to the receiving environments. Environ Int. 92–93:210–219.

Heydecke A, Andersson B, Holmdahl T, Melhus Å. 2013. Human wound infections caused by Neisseria animaloris and Neisseria zoodegmatis, former CDC Group EF-4a and EF-4b. Infect Ecol Epidemiol. 3:20312.

Hibbing ME, Fuqua C, Parsek MR, Peterson SB. 2010. Bacterial competition: Surviving and thriving in the microbial jungle. Nat Rev Microbiol. 8:15–25.

Hughes VM, Datta N. 1983. Conjugative plasmids in bacteria of the “pre-antibiotic” era. Nature. 302:725–726.

Huttenhower C, Gevers D, Knight R, Abubucker S, Badger JH, Chinwalla AT, Creasy HH, Earl AM, Fitzgerald MG, Fulton RS, et al. 2012. Structure, function and diversity of the healthy human microbiome. Nature. 486:207–214.

Jerse AE, Sharma ND, Simms AN, Crow ET, Snyder LA, Shafer WM. 2003. A gonococcal efflux pump system enhances bacterial survival in a female mouse model of genital tract infection. Infect Immun. 71:5576–5582.

Jia B, Raphenya AR, Alcock B, Waglechner N, Guo P, Tsang KK, Lago BA, Dave BM, Pereira S, Sharma AN, et al. 2017. CARD 2017:expansion and model-centric curation of the comprehensive antibiotic resistance database. Nucleic Acids Res. 45:D566–D573.

Jijón S, Wetzel A, LeJeune J. 2007. Salmonella enterica isolated from wildlife at two Ohio rehabilitation centers. J Zoo Wildl Med. 38:409–413.

Jin Y, Yip H-K. 2002. Supragingival calculus: formation and control. Crit Rev Oral Biol Med. 13:426– 441.

Johnston ER, Rodriguez-R LM, Luo C, Yuan MM, Wu L, He Z, Schuur EAG, Luo Y, Tiedje JM, Zhou J, et al. 2016. Metagenomics reveals pervasive bacterial populations and reduced community diversity across the Alaska tundra ecosystem. Front Microbiol. 7:1–16.

Jones KE, Patel NG, Levy MA, Storeygard A, Balk D, Gittleman JL, Daszak P. 2008. Global trends in emerging infectious diseases. Nature. 451:990–993.

Key FM, Posth C, Krause J, Herbig A, Bos KI. 2017. Mining Metagenomic Data Sets for Ancient DNA: Recommended Protocols for Authentication. Trends Genet. 33:508–520.

Khan FA, Söderquist B, Jass J. 2019. Prevalence and diversity of antibiotic resistance genes in Swedish aquatic environments impacted by household and hospital wastewater. Front Microbiol. 10:688.

Knapp CW, Dolfing J, Ehlert PAI, Graham DW. 2010. Evidence of increasing antibiotic resistance gene abundances in archived soils since 1940. Environ Sci Technol. 44:580–587.

Knights D, Kuczynski J, Charlson ES, Zaneveld J, Mozer MC, Collman RG, Bushman FD, Knight R, Kelley ST. 2011. Bayesian community-wide culture-independent microbial source tracking. Nat Methods. 8:761–763.

Könönen E, Wade WG. 2015. Actinomyces and related organisms in human infections. Clin Microbiol Rev. 28:419–442.

Kozak GK, Boerlin P, Janecko N, Reid-Smith RJ, Jardine C. 2009. Antimicrobial resistance in Escherichia coli isolates from swine and wild small mammals in the proximity of swine farms and in natural environments in Ontario, Canada. Appl Environ Microbiol. 75:559–566.

Landecker H. 2016. Antibiotic resistance and the biology of history. Body Soc. 22:19–52.

Landers TF, Cohen B, Wittum TE, Larson EL. 2012. A review of antibiotic use in food animals: Perspective, policy, and potential. Public Health Rep. 127:4–22.

Leatherbarrow AJH, Griffiths R, Hart CA, Kemp R, Williams NJ, Diggle PJ, Wright EJ, Sutherst J, Houghton P, French NP. 2007. Campylobacter lari: Genotype and antibiotic resistance of isolates from cattle, wildlife and water in an area of mixed dairy farmland in the United Kingdom. Environ Microbiol. 9:1772–1779.

Levy SB, Marshall B. 2004. Antibacterial resistance worldwide: Causes, challenges and responses. Nat Med. 10:S122–S129.

Li H. 2013. Aligning sequence reads, clone sequences and assembly contigs with BWA-MEM. arXiv Prepr.:1303.3997.

Li H, Durbin R. 2009. Fast and accurate short read alignment with Burrows-Wheeler transform. Bioinformatics. 25:1754–1760.

Li H, Handsaker B, Wysoker A, Fennell T, Ruan J, Homer N, Marth G, Abecasis G, Durbin R, 1000 Genome Project Data Processing Subgroup. 2009. The Sequence Alignment/Map format and SAMtools. Bioinformatics. 25:2078–2079.

Literak I, Dolejska M, Radimersky T, Klimes J, Friedman M, Aarestrup FM, Hasman H, Cizek A. 2010. Antimicrobial-resistant faecal Escherichia coli in wild mammals in central Europe: Multiresistant Escherichia coli producing extended-spectrum beta-lactamases in wild boars. J Appl Microbiol. 108:1702–1711.

Literak I, Dolejska M, Rybarikova J, Cizek A, Strejckova P, Vyskocilova M, Friedman M, Klimes J. 2009. Highly variable patterns of antimicrobial resistance in commensal Escherichia coli isolates from pigs, sympatric rodents, and flies. Microb Drug Resist. 15:229–237.

Lloyd-price J, Mahurkar A, Rahnavard G, Crabtree J, Orvis J, Brantley Hall A, Brady A, Creasy HH, McCracken C, Giglio MG, et al. 2018. Strains, functions, and dynamics in the expanded Human Microbiome Project. Nature. 550:61–66.

Lu J, Breitwieser FP, Thielen P, Salzberg SL. 2017. Bracken: estimating species abundance in metagenomics data. PeerJ Comput Sci. 3:e104.

Luque I, Echeita A, León J, Herrera-León S, Tarradas C, González-Sanz R, Huerta B, Astorga RJ. 2009. Salmonella Indiana as a cause of abortion in ewes: Genetic diversity and resistance patterns. Vet Microbiol. 134:396–399.

Madden TL, Camacho C, Ma N, Coulouris G, Avagyan V, Bealer K, Papadopoulos J. 2009. BLAST+: architecture and applications. BMC Bioinformatics. 10:421.

Mann AE, Sabin S, Ziesemer K, Vågene ÅJ, Schroeder H, Ozga AT, Sankaranarayanan K, Hofman CA, Fellows Yates JA, Salazar-García DC, et al. 2018. Differential preservation of endogenous human and microbial DNA in dental calculus and dentin. Sci Rep. 8:9822.

Mariano V, McCrindle CME, Cenci-Goga B, Picard JA. 2009. Case-control study to determine whether river water can spread tetracycline resistance to unexposed impala (aepyceros melampus) in Kruger National Park (South Africa). Appl Environ Microbiol. 75:113–118.

Martin JF, Liras P. 1989. Organization and expression of genes involved in the biosynthesis of antibiotics and other secondary metabolites. Annu Rev Microbiol. 43:173–206.

McCann CM, Christgen B, Roberts JA, Su JQ, Arnold KE, Gray ND, Zhu YG, Graham DW. 2019. Understanding drivers of antibiotic resistance genes in High Arctic soil ecosystems. Environ Int. 125:497–504.

Meyer M, Kircher M. 2010. Illumina sequencing library preparation for highly multiplexed target capture and sequencing. Cold Spring Harb Protoc. 2010:pdb.prot5448.

Miller R V., Gammon K, Day MJ. 2009. Antibiotic resistance among bacteria isolated from seawater and penguin fecal samples collected near Palmer Station, Antarctica. Can J Microbiol. 55:37–45.

Mo SS, Urdahl AM, Madslien K, Sunde M, Nesse LL, Slettemeås JS, Norström M. 2018. What does the fox say? Monitoring antimicrobial resistance in the environment using wild red foxes as an indicator. PLoS One. 13:e0198019.

Murray AK, Zhang L, Yin X, Zhang T, Buckling A, Snape J, Gaze WH. 2018. Novel Insights into Selection for Antibiotic Resistance in Complex Microbial Communities. MBio. 9:e00969–18.

Navarro-Gonzalez N, Mentaberre G, Porrero CM, Serrano E, Mateos A, López-Martín JM, Lavín S, Domínguez L. 2012. Effect of Cattle on Salmonella Carriage, Diversity and Antimicrobial Resistance in Free-Ranging Wild Boar (Sus scrofa) in Northeastern Spain. PLoS One. 7:e51614.

Naylor NR, Atun R, Zhu N, Kulasabanathan K, Silva S, Chatterjee A, Knight GM, Robotham J V. 2018. Estimating the burden of antimicrobial resistance: a systematic literature review. Antimicrob Resist Infect Control. 7:58.

Nesme J, Cécillon S, Delmont TO, Monier JM, Vogel TM, Simonet P. 2014. Large-scale metagenomic-based study of antibiotic resistance in the environment. Curr Biol. 24:1096–1100.

Nhung NT, Cuong N V., Campbell J, Hoa NT, Bryant JE, Truc VNT, Kiet BT, Jombart T, Trung N V., Hien VB, et al. 2015. High levels of antimicrobial resistance among Escherichia Coli isolates from livestock farms and synanthropic rats and shrews in the Mekong Delta of Vietnam. Appl Environ Microbiol. 81:812–820.

O’Neill (chair) J. 2014. Antimicrobial Resistance: Tackling a crisis for the health and wealth of nations.

Oh J, Byrd AL, Deming C, Conlan S, Barnabas B, Blakesley R, Bouffard G, Brooks S, Coleman H, Dekhtyar M, et al. 2014. Biogeography and individuality shape function in the human skin metagenome. Nature. 514:59–64.

Oksanen AJ, Blanchet FG, Friendly M, Kindt R, Legendre P, Mcglinn D, Minchin PR, Hara RBO, Simpson GL, Solymos P, et al. 2018. Vegan: Community ecology package. R package version 2.5-3. https://CRAN.R-project.org/package=vegan.

Okubo T, Ae R, Noda J, Iizuka Y, Usui M, Tamura Y. 2019. Detection of the sul2–strA–strB gene cluster in an ice core from Dome Fuji Station, East Antarctica. J Glob Antimicrob Resist. 17:72–78.

Perron GG, Whyte L, Turnbaugh PJ, Goordial J, Hanage WP, Dantas G, Desai MM. 2015. Functional characterization of bacteria isolated from ancient arctic soil exposes diverse resistance mechanisms to modern antibiotics. PLoS One. 10:e0069533.

Poole K. 2005. Efflux-mediated antimicrobial resistance.

Pruden A, Arabi M, Storteboom HN. 2012. Correlation between upstream human activities and riverine antibiotic resistance genes. Environ Sci Technol. 46:11541–11549.

QGIS.org. 2020. QGIS Geographic Information System.QGIS Associationhttp://www.qgis.org.

Quinlan AR, Hall IM. 2010. BEDTools: A flexible suite of utilities for comparing genomic features. Bioinformatics. 26:841–842.

R Core Team. 2020. R: A Language and Environment for Statistical Computing.R Foundation for Statistical Computing,Vienna, Austria.https://www.r-project.org/.

Radhouani H, Igrejas G, Gonçalves A, Pacheco R, Monteiro R, Sargo R, Brito F, Torres C, Poeta P. 2013. Antimicrobial resistance and virulence genes in Escherichia coli and enterococci from red foxes (Vulpes vulpes). Anaerobe. 23:82–86.

Rifkin RF, Vikram S, Ramond JB, Rey-Iglesia A, Brand TB, Porraz G, Val A, Hall G, Woodborne S, Le Bailly M, et al. 2020. Multi-proxy analyses of a mid-15th century Middle Iron Age Bantu-speaker palaeo-faecal specimen elucidates the configuration of the “ancestral” sub-Saharan African intestinal microbiome. Microbiome. 8:62.

Rohland N, Harney E, Mallick S, Nordenfelt S, Reich D. 2015. Partial uracil-DNA-glycosylase treatment for screening of ancient DNA. Philos Trans R Soc B Biol Sci. 370:20130624.

RStudio Team. 2020. RStudio: Integrated Development Environment for R.RStudio, PBC,Boston, MA.http://www.rstudio.com/.

Salter SJ, Cox MJ, Turek EM, Calus ST, Cookson WO, Moffatt MF, Turner P, Parkhill J, Loman NJ, Walker AW. 2014. Reagent and laboratory contamination can critically impact sequence-based microbiome analyses. BMC Biol. 12:87.

Salyers AA, Gupta A, Wang Y. 2004. Human intestinal bacteria as reservoirs for antibiotic resistance genes. Trends Microbiol. 12:412–416.

Schmieder R, Edwards R. 2011. Quality control and preprocessing of metagenomic datasets. Bioinformatics. 27:863–864.

Schneider VA, Graves-Lindsay T, Howe K, Bouk N, Chen H-C, Kitts PA, Murphy TD, Pruitt KD, Thibaud-Nissen F, Albracht D, et al. 2017. Evaluation of GRCh38 and de novo haploid genome assemblies demonstrates the enduring quality of the reference assembly. Genome Res. 27:849–864.

Schubert M, Lindgreen S, Orlando L. 2016. AdapterRemoval v2: Rapid adapter trimming, identification, and read merging. BMC Res Notes. 9:88.

Sharkey LKR, Edwards TA, O’Neill AJ. 2016. ABC-F proteins mediate antibiotic resistance through ribosomal protection. MBio. 7:e01975–15.

Sjölund M, Bonnedahl J, Hernandez J, Bengtsson S, Cederbrant G, Pinhassi J, Kahlmeter G, Olsen B. 2008. Dissemination of multidrug-resistant bacteria into the arctic. Emerg Infect Dis. 14:70–72.

Skurnik D, Ruimy R, Andremont A, Amorin C, Rouquet P, Picard B, Denamur E. 2006. Effect of human vicinity on antimicrobial resistance and integrons in animal faecal Escherichia coli. J Antimicrob Chemother. 57:1215–1219.

Smith DH. 1967. R factor infection of Escherichia coli lyophilized in 1946. J Bacteriol. 94:2071–2072.

Støen OG, Ordiz A, Sahlén V, Arnemo JM, Saebo S, Mattsing G, Kristofferson M, Brunberg S, Kindberg J, Swenson JE. 2018. Brown bear (Ursus arctos) attacks resulting in human casualties in Scandinavia 1977–2016; management implications and recommendations. PLoS One. 13:e0196876.

Subbiah M, Caudell MA, Mair C, Davis MA, Matthews L, Quinlan RJ, Quinlan MB, Lyimo B, Buza J, Keyyu J, et al. 2020. Antimicrobial resistant enteric bacteria are widely distributed amongst people, animals and the environment in Tanzania. Nat Commun. 11:228.

Sugimoto Y, Camacho FR, Wang S, Chankhamjon P, Odabas A, Biswas A, Jeffrey PD, Donia MS. 2019. A metagenomic strategy for harnessing the chemical repertoire of the human microbiome. Science (80-). 366:eaax9176.

Swenson JE, Wabakken P, Sandegren F, Bjärvall A, Franzén R, Söderberg A. 1995. The near extinction and recovery of brown bears in Scandinavia in relation to the bear management policies of Norway and Sweden. Wildlife Biol. 1:11–25.

Tahlan K, Ahn SK, Sing A, Bodnaruk TD, Willems AR, Davidson AR, Nodwell JR. 2007. Initiation of actinorhodin export in Streptomyces coelicolor. Mol Microbiol. 63:951–961.

Tang KL, Caffrey NP, Nóbrega DB, Cork SC, Ronksley PE, Barkema HW, Polachek AJ, Ganshorn H, Sharma N, Kellner JD, et al. 2017. Restricting the use of antibiotics in food-producing animals and its associations with antibiotic resistance in food-producing animals and human beings: a systematic review and meta-analysis. Lancet Planet Heal. 1:e316–e327.

Taylor GA, Kirk H, Coombe L, Jackman SD, Chu J, Tse K, Cheng D, Chuah E, Pandoh P, Carlsen R, et al. 2018. The genome of the North American brown bear or grizzly: Ursus arctos ssp. horribilis. Genes (Basel). 9:598.

Taylor NGH, Verner-Jeffreys DW, Baker-Austin C. 2011. Aquatic systems: Maintaining, mixing and mobilising antimicrobial resistance? Trends Ecol Evol. 26:278–284.

The Center for Disease Dynamics Economics & Policy. 2018. ResistanceMap: Antibiotic use in Sweden.https://resistancemap.cddep.org/CountryPage.php?countryId=35&country=Sweden.

Thomas CM, Nielsen KM. 2005. Mechanisms of, and barriers to, horizontal gene transfer between bacteria. Nat Rev Microbiol. 3:711–721.

Thorpe KE, Joski P, Johnston KJ. 2018. Antibiotic-resistant infection treatment costs have doubled since 2002, now exceeding $2 billion annually. Health Aff. 37:662–669.

Tsukayama P, Boolchandani M, Patel S, Pehrsson EC, Gibson MK, Chiou KL, Jolly CJ, Rogers J, Phillips-Conroy JE, Dantas G. 2018. Characterization of Wild and Captive Baboon Gut Microbiota and Their Antibiotic Resistomes. mSystems. 3:e00016–18.

van der Valk T, Vezzi F, Ormestad M, Dalen L, Guschanski K. 2019. Index hopping on the Illumina HiseqX platform and its consequences for ancient DNA studies. Mol Ecol Resour.

Velsko IM, Frantz LAF, Herbig A, Larson G, Warinner C. 2018. Selection of appropriate metagenome taxonomic classifiers for ancient microbiome research. mSystems. 3:e00080–18.

Venter O, Sanderson EW, Magrach A, Allan JR, Beher J, Jones KR, Possingham HP, Laurance WF, Wood P, Fekete BM, et al. 2016. Global terrestrial Human Footprint maps for 1993 and 2009. Sci Data. 3:160067.

Vittecoq M, Godreuil S, Prugnolle F, Durand P, Brazier L, Renaud N, Arnal A, Aberkane S, Jean-Pierre H, Gauthier-Clerc M, et al. 2016. REVIEW: Antimicrobial resistance in wildlife. J Appl Ecol. 53:519– 529.

Vogwill T, Maclean RC. 2015. The genetic basis of the fitness costs of antimicrobial resistance: A meta-analysis approach. Evol Appl. 8:284–295.

Wallace CC, Yund PO, Ford TE, Matassa KA, Bass AL. 2013. Increase in antimicrobial resistance in bacteria isolated from stranded marine mammals of the Northwest Atlantic. Ecohealth. 10:201–210.

Warinner C, Rodrigues JFM, Vyas R, Trachsel C, Shved N, Grossmann J, Radini A, Hancock Y, Tito RY, Fiddyment S, et al. 2014. Pathogens and host immunity in the ancient human oral cavity. Nat Genet. 46:336–344.

Warinner C, Speller C, Collins MJ, Lewis CM. 2015. Ancient human microbiomes. J Hum Evol. 79:125– 136.

Warner DM, Shafer WM, Jerse AE. 2008. Clinically relevant mutations that cause derepression of the Neisseria gonorrhoeae MtrC-MtrD-MtrE Efflux pump system confer different levels of antimicrobial resistance and in vivo fitness. Mol Microbiol. 70:462–478.

Wickman K. 1969. Kungl. Maj.ts proposition till riksdagen angående förvärv av aktier i AB Kabi och Apoteksvarucentralen Vitrum Apotekare AB. Stockholm, Sweden: Sveriges Riksdag.

Wierup M. 2001. The Swedish experience of the 1986 year ban of antimicrobial growth promoters, with special reference to animal health, disease prevention, productivity, and usage of antimicrobials. Microb Drug Resist. 7:183–190.

Wood DE, Salzberg SL. 2014. Kraken: ultrafast metagenomic sequence classification using exact alignments. Genome Biol. 15:R46.

Zhu YG, Zhao Y, Li B, Huang CL, Zhang SY, Yu S, Chen YS, Zhang T, Gillings MR, Su JQ. 2017. Continental-scale pollution of estuaries with antibiotic resistance genes. Nat Microbiol. 2:16270.

Zottola T, Montagnaro S, Magnapera C, Sasso S, De Martino L, Bragagnolo A, D’Amici L, Condoleo R, Pisanelli G, Iovane G, et al. 2013. Prevalence and antimicrobial susceptibility of Salmonella in European wild boar (Sus scrofa); Latium Region - Italy. Comp Immunol Microbiol Infect Dis. 36:161– 168.

